# COX2-independent and COX2-dependent effects of naproxen on bone quality, osteocytes, and fatigue fracture healing in male and female mice

**DOI:** 10.1101/2025.08.20.669939

**Authors:** Alexandra Ciuciu, Eric McLaughlin, Adriana Ciuciu, Kelly C. Barrientos, Trenton LaPoint, Ryan E. Tomlinson

## Abstract

Regular non-steroidal anti-inflammatory drug (NSAID) use increases stress fracture risk, but the mechanisms remain unclear. Here, we used Ptgs2-Y385F mice, which lack cyclooxygenase 2 (COX2) enzyme activity, to test the hypothesis that naproxen decreases strain adaptive bone remodeling in a COX2-dependent manner and decreases bone toughness and fracture resistance through COX2-independent effects. MicroCT and mechanical testing showed minimal baseline differences between Ptgs2-Y385F and wild-type (WT) mice. Following non-damaging forelimb compression, naproxen decreased load-induced bone formation in WT, but not Ptgs2-Y385F mice, consistent with a COX2-dependent effect. In contrast, naproxen reduced bone toughness and post-yield deformation across genotypes and doses, supporting a COX2-independent mechanism. Histologically, naproxen increased empty osteocyte lacunae in both genotypes, while osteoblast number, total lacunae, and perilacunar labeling were increased only in WT mice. Naproxen pre-treatment before fatigue fracture produced mild, sex-specific effects on fracture initiation and healing. Analysis of osteocyte dendritic networks in uninjured femurs revealed sexually dimorphic changes in dendrite number and density in naproxen-treated mice as compared to vehicle. In total, naproxen influences bone through both COX2-dependent and COX2-independent mechanisms, with some sexual dimorphism. These findings confirm that regular NSAID usage compromises skeletal health and underscore the need for new pain management strategies.

## INTRODUCTION

Opioids and non-steroidal anti-inflammatory drugs (NSAIDs) are the most common and effective pharmaceutical pain management option. Given the increased risks of abuse, side effects, and healthcare costs of opioids, NSAIDs are often preferred^1,2^. Active individuals commonly use NSAIDs to manage musculoskeletal pain from exertion and prophylactically to extend physical activity periods^3,4^. Increasing evidence from human and animal studies supports a correlation between regular NSAID use and increase skeletal fatigue injuries, termed stress fractures^5–8^. Stress fractures form from repetitive physical strains that introduce microcracks into bone that coalesce into a fracture^9–11^. Symptoms of an early stress fracture (e.g. pain and inflammation) may be attributed to muscle soreness and lead to self-medication with NSAIDs, leading to delayed or missed stress fracture diagnoses^10,11^. Despite relieving fracture-related pain effectively, NSAID use is linked to reduced strain-adaptive bone remodeling and bone toughness, elevating stress fracture risk. The underlying mechanisms driving these effects are incompletely understood.

The cyclooxygenase enzymes, COX1 and COX2, convert arachidonic acid to prostaglandins that can initiate pain, fever, and inflammation. NSAIDs inhibit this conversion by either selectively inhibiting one enzyme or non-selectively inhibiting both enzymes^12^. COX1 is constitutively active, whereas COX2 is considered inducible in the cases of disease, inflammation, or injury, but recent evidence supports a critical role for COX2 in cardiac and reproductive systems^13–15^. Consequently, both non-selective and selective NSAIDs may cause off-target effects, some of which occur independently of COX signaling entirely^16–22^. Further, on-target NSAID inhibition of the cyclooxygenase pathway can also be detrimental in the context of bone. Prostaglandins, particularly Prostaglandin E2 (PGE2), influence both bone formation and resorption, and PGE2 is also involved in bone mechanosensation^23,24^. Additionally, the release of prostaglandins during bone injury triggers pain and stimulates cytokines and growth factors required for fracture healing; thus, NSAID use can delay or diminish fracture healing^7,8,25,26^.

A recent study in U.S. army soldiers revealed that using any NSAID regularly increased stress fracture incidence, with naproxen use correlating to the largest increase^6^. Similarly, we have observed this same correlation between regular NSAID use and increased stress fracture incidence in the general population^5^. Other pre-clinical and clinical studies report conflicting effects of regular NSAID use on bone’s response to mechanical forces and fracture healing^7,8,25,27–32^. Differences in results may be caused by variations in dosage, drug delivery method, sex, study design, and NSAID chemical structures. In fact, we previously reported that naproxen, but not aspirin, administered to female C57BL/6J mice in drinking water for 15 days significantly decreased load-induced bone formation, diminished bone toughness, and altered collagen fibril thickness^7^. Although both naproxen and aspirin are non-selective NSAIDs and inhibit both COX1 and COX2, they differ substantially in their mode and duration of COX inhibition. Additionally, previous studies support that naproxen has significant effects in bone and other tissues^6,33–36^.

This study used Ptgs2-Y385F mice, harboring a point mutation that inhibits COX2 cyclooxygenase activity similarly to the inhibition of a COX2-selective NSAID, to assess effects of naproxen, a non-selective NSAID, on bone that are COX2-dependent or -independent. We investigated the effects of naproxen on bone composition and quality when compared to vehicle treatment. Special attention was given to osteocytes, bone cells regulating mechanosensation and transduction, osteoblast and osteoclast driven bone remodeling, perilacunar canalicular remodeling (PLR), and paracrine and endocrine signaling^38–48^. We hypothesized that naproxen increases fatigue fracture risk by diminishing strain adaptive bone remodeling through COX2-dependent mechanisms, but also decreasing bone toughness through COX2-independent mechanisms that influence osteocytes and their dendritic networks. These findings improve current understanding of skeletal effects of NSAIDs, highlight potential off-target effects of naproxen increasing fatigue fracture risk, and reveal sexual dimorphism in the response of bone to regular NSAID use.

## METHODS

### Mice

The Institutional Animal Care and Use Committee (IACUC) of Thomas Jefferson University (Protocol #01919) approved all experiments involving mice. Two Ptgs2-Y385F heterozygous breeding pairs were purchased from the Jackson Laboratory (Strain #008101) and used to breed all further mice in the Ptgs2-Y385F lineage. Breeding schemes consisted of either heterozygous pairings or homozygous males paired with heterozygous females to increase homozygous offspring ratios. Mice homozygous for the Ptgs2-Y385F mutation were compared to their wild-type (WT) or heterozygous littermates unless stated otherwise. Mice were housed in groups of up to five while receiving standard rodent chow (LabDiet 5001). Mice were aged until skeletally mature (14-16 weeks) before experimental start and both sexes of mice were used. Additionally, a cohort of 20 C57BL/6J females were purchased from the Jackson Laboratory (Strain #000664) at 14 weeks old and allowed to acclimate to the animal facility until 16 weeks old while being housed in groups of five. Finally, femurs were harvested from a cohort of 20-week-old and 52-week-old male WT littermates from our colony (C57BL/6J background) to compare to Ptgs2-Y385F homozygous males of the same ages and monitor age-related effects of the Ptgs2-Y385F mutation.

### NSAID Dosing

Treatment groups included vehicle (ddH_2_O), low dose naproxen sodium (10.9 mg/kg/day), or high dose naproxen sodium (41.6 mg/kg/day). Mice were allocated to each treatment group based on their genotype to reach a goal sample size defined for each experiment. Investigators were not blinded to animal genotype or treatment during allocation or animal handling. Naproxen sodium or vehicle was delivered as drinking water and replenished every two to three days. Doses were calculated with an assumed consumption of 4 mL of drinking water per day per mouse with an estimated average mouse mass of 25 grams. High and low doses were adapted from comparable human doses to mice using allometric scaling provided by the FDA Center for Drug Evaluation and Research^49^. The daily recommended dose of naproxen sodium for musculoskeletal pain relief in humans is 3.38 mg/kg, which translates to 41.6 mg/kg in mice. Low dose naproxen was calculated to be about 25% of the daily recommended dose, 10.9 mg/kg/mouse, and high dose naproxen was calculated to be 100% of the daily recommended so, 41.6 mg/kg/mouse. Neither dose was higher than the maximum human recommended dose of 10.2 mg/kg, or 125.5 mg/kg in mice. Vehicle or naproxen sodium drinking water was administered 24 hours before experimental start. All genotype characterization and aging experiments used mice receiving only vehicle drinking water. The non-damaging forelimb loading cohort received treatment for 15 total days while the fatigue fracture induction cohort had a treatment period of 30 total days. All C57BL/6J females received treatment for 30 days, with 10 mice being given low dose naproxen sodium and 10 mice being given high dose naproxen sodium. Low dose naproxen sodium will be referred to as naproxen sodium or naproxen, with high dose naproxen sodium groups labeled as “high dose naproxen”. During endpoint measurements involving image analysis, investigators were blinded to animal genotype and treatment.

### Mechanical Loading

Mice were mechanically loaded using axial forelimb compression to either induce lamellar bone formation or fatigue fracture at the ulnar mid-diaphysis, as previously established^7,50^. Before experimental start, mice were intraperitoneally injected with 0.12 mg/kg buprenorphine, and mice were anesthetized throughout experimentation using isoflurane (2-3%) inhalation. After confirming deep anesthesia through hindpaw pinch, the mouse’s right forelimb was secured in a custom fixture for loading in a material test system (TA Instruments Electroforce 3200 Series III) that recorded force and displacement digitally. Before experiments began, mice received a 0.3 N compressive preload to secure their forelimbs in place.

In lamellar bone formation experiments, mice received either vehicle or naproxen sodium (10.9 mg/kg) drinking water for 15 total days. During treatment, mice were subjected to 100 cycles of a 2 Hz rest-inserted sinusoidal waveform with a peak force of 3.0 N. This loading was repeated three times per week over two weeks for a total of six loading bouts. As shown in previous publications, a peak force of 3.0 N generates an approximate average peak compressive strain of 3000 µɛ at the ulnar mid-diaphysis, inducing bone formation without causing damage^51,52^. Mice were returned to their cages with unrestricted activity after each loading bout and tissues were harvested on day 15.

In fatigue fracture experiments, treatment cohorts were divided between vehicle or naproxen sodium (10.9 mg/kg) and were pre-treated for 15 days before fracture induction. Loading parameters were chosen as a function of ultimate force and displacement to fatigue fracture^53^. First, previously sacrificed heterozygous mice (n=3-4 per sex) outside of the treatment groups were used to determine average ultimate force for males and females separately by having their left forelimb monotonically compressed to failure by a displacement ramp program (0.1 mm/s). Second, displacement to fatigue fracture formation was determined using the same sacrificed mice by subjecting their right forelimbs to a sinusoidal waveform at 2 Hz and 75% of the ultimate force until complete fatigue fracture. A heat lamp was used during fatigue fracture cycling to minimize mechanical differences in forelimbs due to rigor mortis. For fatigue fracture cohorts, 3.3 N was determined to be 75% of the ultimate force to complete forelimb failure. Once an average ultimate displacement per sex was determined, 75% of the total displacement was calculated to be used on treatment cohorts to generate a fatigue fracture at the ulnar mid-diaphysis. The targeted displacement was 0.865 mm for male mice and 0.691 mm for female mice, regardless of treatment or genotype. Thus, for fatigue fracture experiments, mice received a compressive preload of 0.3 N to fix their forelimbs in place, followed by a cyclic sinusoidal waveform with a peak force of 3.3 N at a frequency of 2 Hz until a total displacement of 0.865 mm for males and 0.691 mm for females, relative to the displacement measured on the 10^th^ cycle of compression. This protocol has been demonstrated to generate a non-displaced fatigue fracture 1 mm distal to the ulnar mid-diaphysis and results in woven bone formation^54^. After fatigue fracture induction, mice were returned to their cages with unrestricted cage activity while continuing to receive vehicle or naproxen sodium drinking water, and tissues were harvested on day 30.

### MicroCT

In all experiments, unfixed, non-loaded femurs were harvested, cleaned of soft tissue, and stored in gauze soaked in phosphate-buffered saline (PBS) at -20°C until microCT scanning. Scanning was done using a Bruker Skyscan 1275 microCT system with a 1 mm aluminum filter. Femurs were scanned using 55 kV and 181 µA with a 74 ms exposure time and a 13 µm isometric voxel size. Transverse plane scan slices were obtained by placing the femur parallel to the z-axis of the scanner. Reconstruction and analysis were done using NRecon and CTAn (Bruker), respectively. Trabecular bone geometry was analyzed across a 1 mm region of the femoral metaphysis starting at 0.5 mm below the growth plate and cortical bone geometry was analyzed across a 1 mm region at the femoral mid-diaphysis. Measured outcomes were bone mass, density, and geometry.

For fatigue fracture experiments, unfixed loaded (injured) and non-loaded (intact) forelimbs were harvested 15 days after injury without cleaning of soft tissue and stored in PBS-soaked gauze at -20°C until microCT scanning. Forelimbs were scanned using 55 kV and 181 µA with a 74 ms exposure time and a 10 µm isometric voxel size. Transverse plane scan slices were obtained by placing the forelimb parallel to the z-axis of the scanner. Reconstruction and analysis were performed as described above. Forelimbs bone calluses and radial and ulnar geometry were analyzed. Bone callus analysis had measured outcomes of woven bone volume, extent, and bone mineral density, crack size, and crack location relative to limb length. After scanning, both loaded (injured) and non-loaded (intact) forelimbs were subjected to mechanical testing by monotonic compression until failure, whereafter the non-loaded forelimbs were scanned again to analyze crack formation during compression in the non-loaded forelimb. Radial and ulnar geometry were analyzed in non-loaded (intact) limbs across 0.5 mm regions at two locations: starting at 6.92 mm from the olecranon process (termed Proximal) or starting at 7.99 mm from the olecranon process (termed Distal). These two regions were determined to be the average locations for crack formation in WT and Ptgs2-Y385F mice respectively, thus the bone geometries of the radius and ulna were examined for potential genotype or NSAID effects on cortical bone geometry in these regions. Reconstruction and analysis were performed as stated above and measured outcomes included bone mass, density, geometry, and the starting point of the fatigue fracture crack.

### Standard Three-Point Bending

Unfixed, non-loaded femurs were mechanically tested using standard three-point bending to measure their structural and material properties. Femurs that were previously scanned using microCT were stored in PBS-soaked gauze at -20°C until testing, then thawed to room temperature. Thawed femurs were oriented on a stationary fixture of a material testing system (TA Instruments Electroforce 3200 Series III) with femoral condyles facing down and a bending span of approximately 7.91 mm. A 0.3 N preload was applied to secure the femur in place then a monotonic displacement ramp of 0.1 mm/s until failure was applied while force and displacement values were collected digitally. The resulting force-displacement curve was converted using microCT-based geometry from the femoral mid-diaphysis slice and a custom GNU Octave script to a stress-strain curve.

### Forelimb Mechanical Testing

MicroCT-scanned loaded (injured) and non-loaded (intact) forelimbs from fatigue fracture experiments were stored in PBS-soaked gauze at -20°C until testing, then thawed to room temperature. Thawed forelimbs were placed into the same fixture of the material testing system (TA Instruments Electroforce 3200 Series III) as was used for fatigue fracture induction. Forelimbs were monotonically compressed to failure at 0.1 mm/s, a protocol previously shown to measure structural properties of woven bone in a healing fracture^55^. During compression, force and displacement data were collected digitally and used to calculate strength, stiffness, and total energy to failure using a custom GNU Octave script. Results were normalized to the contralateral, non-injured limb.

### Dynamic Histomorphometry and Osteocyte Fluorophore Labeling

To quantify lamellar bone formation during loading, calcein (10 mg/kg) and alizarin red (30 mg/kg) were intraperitoneally injected into mice on days 5 and 12 of the experiment. Forelimbs were harvested on day 15, fixed for 24-48 hours in neutral buffered formalin (NBF), dehydrated in either graded ethanol or 100% xylenes, and embedded in polymethylmethacrylate (PMMA). Once embedded, forelimbs were sectioned at the ulnar mid-diaphysis using a low-speed diamond wafering saw (Isomet 1000) at a thickness of about 500 µm, mounted on glass slides using Eukitt mounting medium, and hand-sanded to a thickness of 70-100 µm for imaging and analysis. Images were captured on a Zeiss LSM 800 confocal microscope at 20x and analyzed for periosteal (Ps) and endosteal (Es) mineralizing surface per bone surface (MS/BS), mineral apposition rate (MAR), and bone formation rate (BFR/BS) as defined by the ASBMR committee for Histomorphometry Nomenclature^56^. Relative (r) parameters were calculated as the difference between loaded and non-loaded limbs. The number of samples with single labels or double labels are listed in Supplementary Table 3 and any sample without double labeling on the endosteal or periosteal surfaces were assigned 0.3 µm/day for MAR on that surface.

After imaging for dynamic histomorphometry, a cohort of female WT samples were stained with Vectashield Plus antifade mounting medium with DAPI (H-1500) (Vector Laboratories) for nuclear labeling and forelimb sections were imaged at 40x on a Zeiss LSM 800 confocal microscope. Osteocytes labeled with either calcein, alizarin red, or both fluorophores were quantified as a measure of remodeling in the perilacunar space.

### Picrosirius Red Staining

To quantify changes to fibrillar collagen organization, non-loaded tibias from all treatment groups were harvested, dissected free of soft tissue, fixed for 24-48 hours in NBF, and decalcified in 14% EDTA for 30-35 days.

Tibias from mice in lamellar bone formation experiments (15 days) were then embedded in paraffin and sectioned at 4 µm thickness in cross sections. Tibial sections were deparaffinized in 100% xylenes for 5 minutes, then gradually rehydrated in decreasing alcohol gradients from 100% to 70% ethanol solutions. Samples were then placed under a running tap water bath for 2 minutes and air dried. Afterwards, samples were covered in phosphomolybdic acid for 2 minutes, rinsed with 1X PBS, covered with picrosirius red solution for 60 minutes, dried, and covered in 0.1 N hydrochloric acid for 2 minutes. After staining, samples were dehydrated in increasing alcohol gradients from 70% to 100% ethanol, then submerged in 100% xylenes (Polysciences, #24901). Samples were mounted onto microscope slides using Eukitt mounting solution and imaged with a circularly polarized light microscope (Nikon Eclipse LV100POL) at 20x. Applying this procedure, the birefringence of collagen in bone changes color from shorter to longer wavelengths (from green to red) as collagen fiber thickness increases^57^. Best-fit color thresholds were determined manually by individual pixel categorization into either red (thick), yellow (medium), or green (thin) fibrils in four biological replicates using NIS-Elements AR 4.5.00 (Nikon). Best-fit color thresholds for green, yellow, and red were then applied to all images and the fraction of area in the tibial cross section occupied by each color was measured. Representative images were tiled using FIJI^58,59^. Two technical replicates were stained from each mouse, and one replicate was chosen for analysis as determined by stain consistency under brightfield imaging.

Separately, tibias from mice in fatigue fracture experiments (30 days) were then flushed of bone marrow, sunk in 30% sucrose solution, and embedded in Tissue-Tek O.C.T. Compound and frozen at –20°C before sectioning at 7 µm thickness in cross sections. Samples were rinsed with 1X PBS for 5 minutes three times to remove O.C.T., covered in picrosirius red solution for 60 minutes, dab dried, and covered in 0.1 N hydrochloric acid solution for 2 minutes. Samples were dehydrated and mounted as stated above. Imaging and analysis were done using the same microscope, programs, and settings as for lamellar bone formation samples above. Three technical replicates were stained from each mouse, and one technical replicate was used for analysis as determined by sectioning and stain consistency under brightfield imaging.

### Hematoxylin and Eosin Staining

Non-loaded tibias from mice in lamellar bone formation experiments (15 days) were harvested, fixed in NBF overnight, decalcified for 30 days in 14% EDTA, and embedded in paraffin. Tibias were then sectioned longitudinally at 4 µm onto microscope slides and stained with hematoxylin and eosin at the Thomas Jefferson University Translational Research and Pathology Shared Resource Facility. Samples were imaged on a Nikon E-800 Brightfield Microscope at 4x to visualize growth plates or at 40x to visualize osteoblasts in the metaphysis or osteocytes in cortical bone of the tibial diaphysis. Quantifications of average growth plate thickness, osteoblast number per bone perimeter, and osteocyte lacunae number per bone area were done manually in FIJI^58^.

### Immunofluorescence

Non-loaded tibias from mice in lamellar bone formation experiments (15 days) were harvested, fixed in NBF overnight, decalcified for 30 days in 14% EDTA, and embedded in paraffin. Tibias were sectioned longitudinally at 4 µm onto microscope slides, deparaffinized in 100% xylenes, rehydrated in decreasing gradients of alcohol from 100% to 70% ethanol, and hydrated with 1X PBS. Samples were then submerged in 0.05% citraconic anhydride solution (pH adjusted to 7.4) at 100°C for 30 minutes, followed by gradual cooling while submerged in citraconic anhydride solution in a 4°C fridge for 10 minutes. After antigen retrieval, samples were washed using 1X PBS and 1X Tris-buffered saline with 0.1% Tween 20 detergent (TBST). Samples were then covered in a blocking solution of 10% donkey serum in 1X PBS for one hour at 4°C. The blocking solution was then removed, and samples were covered with a primary antibody for matrix metalloproteinase 13 (rabbit anti-MMP-13 polyclonal antibody) at 4°C overnight in a humidified chamber (Abcam 39012; 1:50). The primary antibody was then washed off using 1X PBS and 1X TBST before samples were incubated with a donkey, anti-rabbit secondary antibody conjugated to AlexaFluor 594 (Jackson ImmunoResearch Labs Inc. #711-585-152, 1:1000) for 1 hour at 4°C in a dark, humidified chamber. Excess antibody was then removed using 1X PBS and samples were mounted using Vectashield Plus antifade mounting medium with DAPI (H-1500) (Vector Laboratories). Samples were imaged on a Zeiss LSM 800 confocal microscope at 40x and captured using Z-stacking at a thickness of 4-4.1 µm thick before being stacked and counted for the number of osteocytes positive for MMP-13 in their perinuclear space per bone area using FIJI^58^.

### Osteocyte Dendritic Network Analysis

Non-loaded femurs from mice in fatigue fracture experiments (30 days) were harvested, fixed in NBF for 24-48 hours, decalcified for 21-30 days in 14% EDTA, sunk in 30% sucrose, and embedded in Tissue-Tek O.C.T. compound at -20°C. Femurs were then sectioned in the femoral diaphysis at a thickness of 20 µm in cross sections. After sectioning, samples were placed in -20°C overnight, thawed to room temperature, and washed with 1X PBS three times for 10 minutes to remove O.C.T. After washing, samples were permeabilized with a 0.5% Triton X-100 solution (made with 3% Bovine Serum Albumin (BSA) in 1X PBS) horizontally for 30 minutes at room temperature. Once permeabilized, samples were washed two times for 5 minutes in 1X PBS. A working solution of Phalloidin conjugated to AlexaFluor 488 (Invitrogen A12379) was made fresh at a concentration of 1:40 of the stock solution diluted in 1X PBS. Samples were covered in this working solution horizontally for 2 hours at room temperature with gentle rocking and blocked from light exposure. After staining, samples were washed two times in 1X PBS to remove excess phalloidin and mounted using Vectashield Plus antifade mounting medium with DAPI (H-1500) (Vector Laboratories). Stained samples were stored at -20°C before use and imaged using a Zeiss LSM 800 confocal microscope within 2-3 days of staining to avoid fluorescent fading. Images were taken of osteocyte dendrites at 63x using Z-stacking with a thickness of 15-15.1 µm. Collected images were separated into a nuclear and a dendritic channel and the tubeness function on FIJI was applied to the dendritic channel (sigma 0.099). This function rates points in the image by how tube-like they are to distinguish dendrites from background noise signal. The channels were then stacked using a maximum intensity projection and the colors were merged with nuclei in blue and dendrites in green. Color channels were adjusted separately using the color balance tool and green was auto adjusted four times while blue was auto adjusted once across all images for better visualization during analysis. Analysis was done using a custom “Moving Band Method” or by manual dendrite counting, and in all cases images were blinded during analysis.

In the “Moving Band Method” the maximum intensity projection of a region was divided in two channels (blue for DAPI and green for phalloidin/dendrites) and then the blue channel was used to create a mask of the nuclei by thresholding the image until all nuclei are selected without holes or distortions. This mask was then despeckled and any non-nuclear selections were removed. The resulting nuclei masks were used to create bands around the nucleus by dilating the original nucleus to a set number of iterations and subtracting smaller iterations using the binary/dilate and image calculator tools in FIJI. The bands analyzed were at 12 iterations minus the original nucleus, 36 iterations minus 24 iterations, 60 iterations minus 48 iterations, and 96 iterations minus 84 iterations. These created bands at 1.2 µm, 2.4-3.6 µm, 4.8-6.0 µm, and 8.4-9.6 µm from the nucleus that represented changing dendritic density as distance from the nucleus increased. Bands were analyzed separately, and dendritic density was estimated by adding the banded image with the original green/phalloidin channel to create a band of dendrites at the region of interest (ROI). Once this image was created, a threshold was applied using the default threshold settings on FIJI and this threshold divided the area of the band that was occupied by signal by the total area of the bands to generate a % ROI that was used to compare dendritic densities across treatment groups.

For manual counting of dendrites, a similar method of creating bands of dendrites was used with initial dendrites (sprouting from the nucleus) being counted at a band of 20 iterations minus the original nucleus mask (a distance of 2.0 µm from the nucleus) and terminal dendrites (farthest from the nucleus) being counted at a band of 100 iterations minus 80 iterations (8.0-10 µm from the nucleus). The dendrites from three osteocytes from separate bone sections per biological replicate were counted and averaged. All image processing and analysis of dendrites was done using FIJI^58^.

### Statistics

Statistical analysis was performed using Prism 10 (GraphPad Software Inc.) and statistical outliers were calculated using either Grubbs’ test or ROUT. Due to varying sample sizes stemming from inconsistent litter sizes and frequencies in the Ptgs2-Y385F mouse line and testing cohorts, unpaired two-tailed t tests with Welch’s correction, Welch one-way ANOVAs, and two-way ANOVAs were used for statistical comparisons as stated with each result. All analyses were performed without assuming equal standard deviations between groups. An adjusted p-value of less than 0.05 was considered significant and adjusted p-values below 0.1 were considered trending. Additionally, in cases where males and females did not display different effects due to sex, sexes were combined for increased statistical power but still delineated as full circles for females and empty circles for males. Finally, all bar graphs, unless stated otherwise, include error bars delineating standard deviation.

## RESULTS

### The Ptgs2-Y385F mutation does not significantly diminish bone geometry and mechanical performance or mouse size

We performed microCT and standard three-point bending analysis of unfixed, non-loaded femurs from Ptgs2-Y385F, heterozygous, and wild-type (WT) littermates of both sexes that received only vehicle drinking water. Trabecular and cortical bone were analyzed using separate regions of interest (**Fig. 1A-C**). Femur lengths between genotypes were not significantly different at 16-18 weeks of age (**Fig. 1D**). Differences between genotypes in trabecular or cortical bone geometry were mild (**Fig. 1E-K**). Changes due to genetic mutation were slightly more prominent in females than males, with Ptgs2-Y385F females exhibiting increases in trabecular BMD, BV/TV, thickness, and number as well as cortical bone area per tissue area and cross-sectional thickness as compared to WT. These findings were confirmed on a separate cohort of mice at 20 weeks of age (**S. Fig. 1A-L and S. Fig. 2A-J**). By three-point bending, Ptgs2-Y385F mice have significant increases in ultimate strain, ultimate bending energy, post-yield energy, and toughness compared to WT littermates at 16-18 weeks (**Fig. 1L-Q**). Analyzing sexes separately showed similar results to combined data, except male Ptgs2-Y385F mice had reduced ultimate stress, while females showed no effects due to genotype. However, these differences were not visible at 20 weeks of age (**S. Fig. 1M-R**). Finally, body masses at 18 weeks were compared between Ptgs2-Y385F and WT mice and no significant differences were seen with sexes combined (**Fig. 1R**), and only a mild decrease was seen in male Ptgs2-Y385F mice compared to WTs when sexes were separated (**S. Fig. 1S-T**). In total, Ptgs2-Y385F mice are comparable to WT littermates at skeletal maturity in bone length, size, and mechanical performance and body mass.

**Figure 1:**
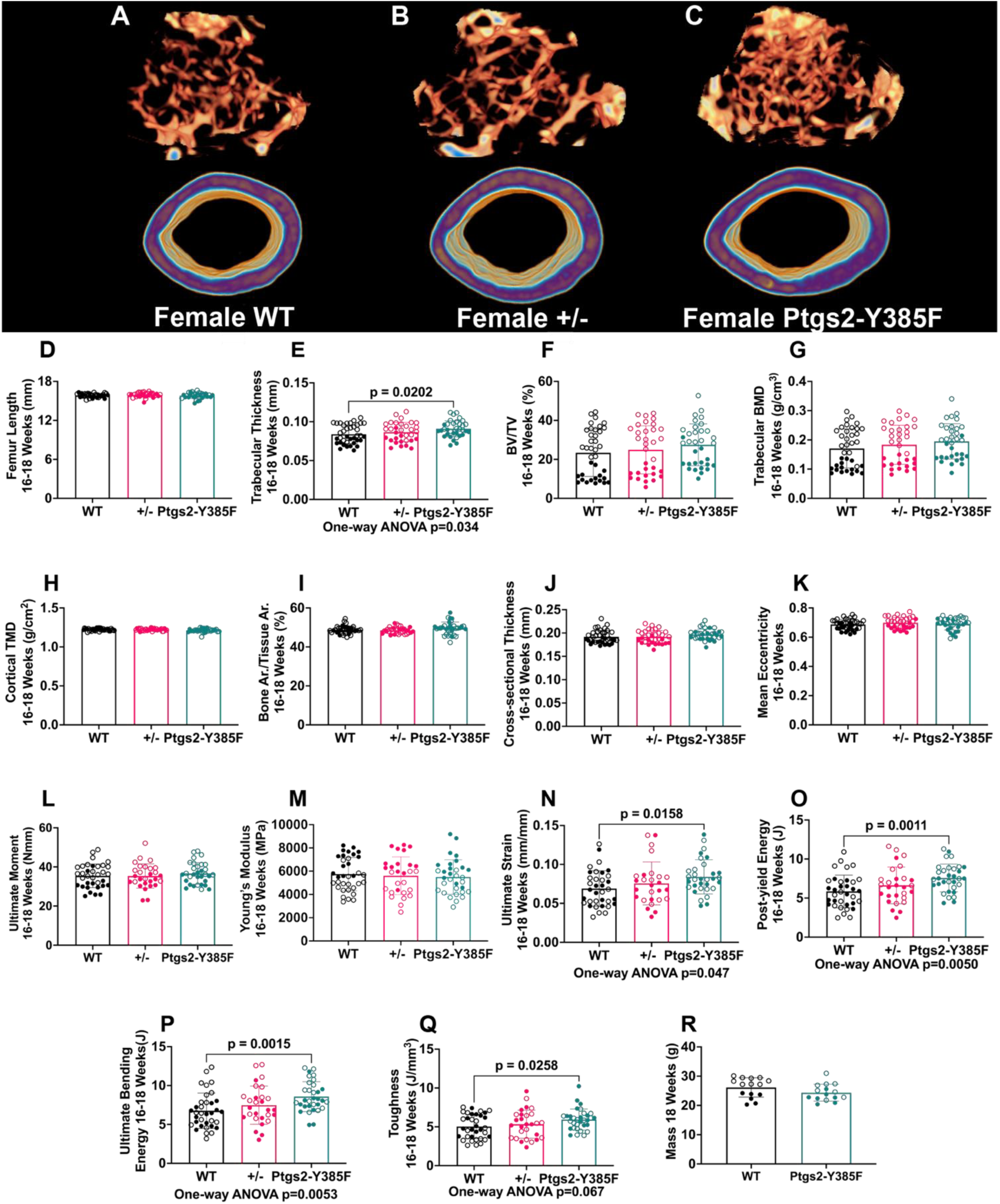
Ptgs2-Y385F mice are comparable to WT littermates in size, bone geometry, and bone mechanical performance at 16-18 weeks old. Bone geometry was quantified using microCT scanning and reconstruction of non-loaded femurs and femoral mechanical performance was measured using standard three-point bending. (A-C) Representative microCT reconstructions of trabecular and cortical bone regions for female wild-type (WT), heterozygous (+/-), and Ptgs2-Y385F mice. Bone geometry parameters include (D) femur length, (E) trabecular thickness, (F) trabecular bone volume per tissue volume (BV/TV), (G) trabecular bone mineral density (BMD), (H) cortical tissue mineral density (TMD), (I) cortical bone area per tissue area (Bone Ar./ Tissue Ar.), (J) cortical cross-sectional thickness, and (K) cortical mean eccentricity. n=31-38 per group. Mechanical performance parameters include (L) ultimate moment, (M) Young’s modulus, (N) ultimate strain, (O) post-yield energy, (P) ultimate bending energy, and (Q) toughness. n=28-35 per group. (R) Body mass at 18 weeks is included as a measure of size at skeletal maturity. n=15 per group. A p value below 0.05 was considered significant and a p value below 0.1 was considered trending. Welch’s ANOVA F statistics for these analyses ranged from 0.4033 to 3.546. Female replicates are displayed as filled circles and male replicates as empty circles.

**Figure 2:**
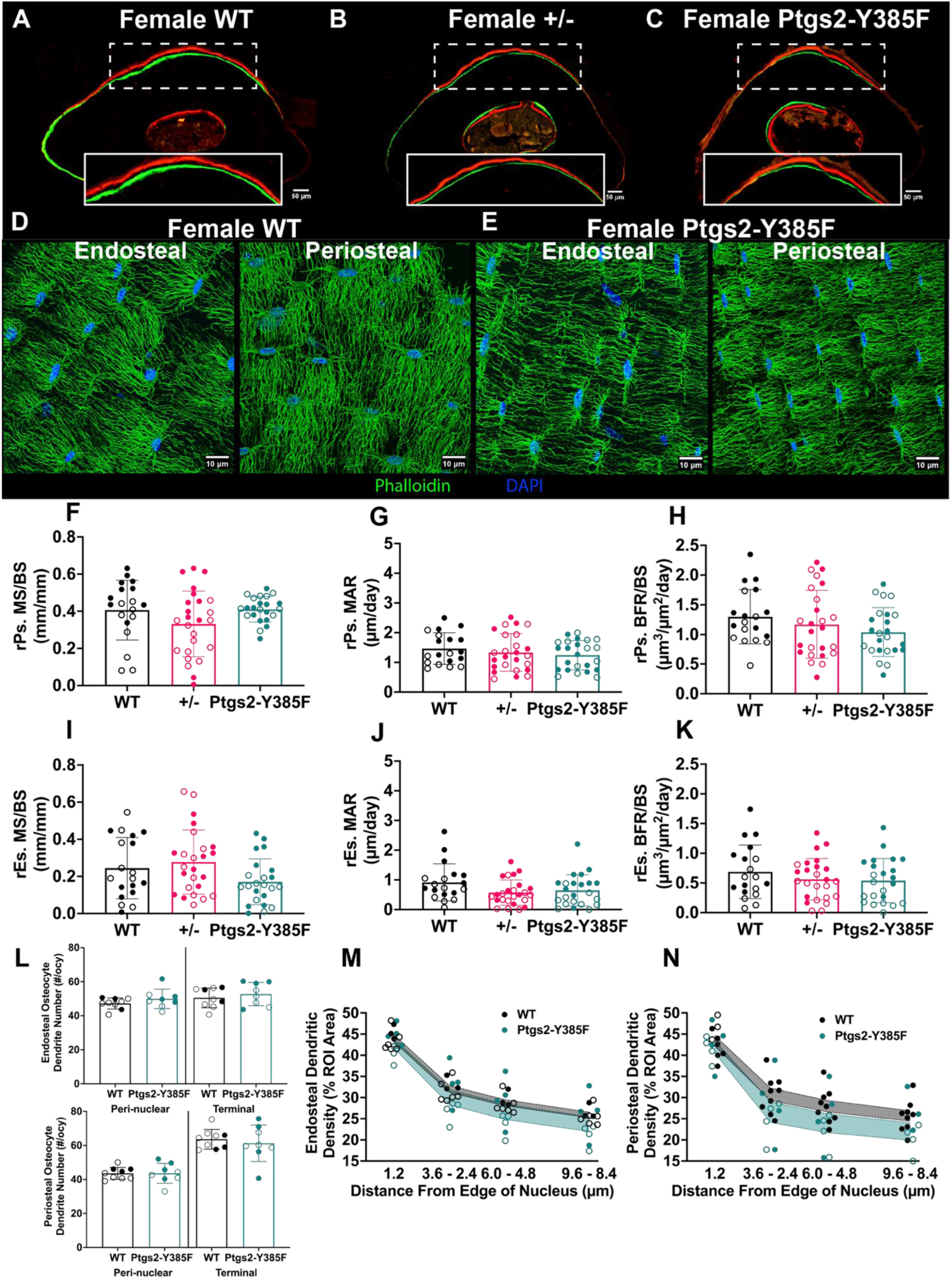
The Ptgs2-Y385F mutation does not inhibit load-induced bone formation or significantly change the osteocyte dendritic network. Bone formation in response to uniaxial forelimb compression was measured in ulnar mid-diaphysis sections using dynamic histomorphometry with calcein (green) and alizarin red (red) bone formation labels in mice aged 16-18 weeks at experiment end. Osteocyte dendritic networks were visualized in mid-cortical sections of non-loaded femurs using phalloidin F-actin staining (green) and DAPI nuclear staining (blue), and networks were quantified using dendrite counting near the nucleus (peri-nuclear) and at a standard distance away from the nucleus where dendrites commonly ended (terminal) and dendritic density was measured using the moving band method in mice aged 20 weeks at experiment end. (A-C) Representative sections from female WT, heterozygous (+/-), and Ptgs2-Y385F mice. (D-E) Representative sections from female WT and Ptgs2-Y385F mice showing the dendritic network near the endosteal and the periosteal surfaces of the bone. Measured outcomes included (F) relative periosteal mineralizing surface per bone surface (rPs. MS/BS), (G) relative periosteal mineral apposition rate (rPs. MAR), (H) relative periosteal bone formation rate per bone surface (rPs. BFR/BS), (I-K) and corresponding endosteal surface parameters. n=19-25 per group. (L) Dendrite number per osteocyte in the peri-nuclear and terminal regions considered. Dendritic density as measured by the moving band method in osteocytes near the (M) endosteal and (N) periosteal surfaces of the bone and the boundaries of the standard error of the mean are colorized alongside individual replicates. n=8-9 per group. A p value below 0.05 was considered significant and a p value below 0.1 was considered trending. Welch’s ANOVA F statistic for dynamic histomorphometry measures ranged from 0.6934 to 3.329. Welch’s t test F statistic for dendrite quantification measures ranged from 1.397 to 3.061. Female replicates are displayed as filled circles and male replicates as empty circles.

Next, Ptgs2-Y385F males at 20 weeks and 52 weeks of age were compared to age-matched C57BL/6J WT male mice from our colony. As before, trabecular and cortical bone were analyzed using separate regions of interest in mice at each age (**S. Fig. 3A-D)**. Ptgs2-Y385F mice had significantly decreased femoral length than WT mice at all ages, but showed only slightly decreased femoral growth over the 32-week period (**S. Fig. 3E**). Trabecular geometry measures between genotypes were also significantly different at both ages, but overall changes showed similar decreases with aging across genotypes (**S. Fig. 3F-H**). Ptgs2-Y385F mice showed minimal age-related changes in bone marrow area and polar moment of inertia between 20 and 52 weeks, in contrast to the significant increases seen in aging WT mice (**S. Fig. 3I-N**). Differences in cortical bone geometry resulted in reduced mechanical performance of Ptgs2-Y385F bone, most notably in ultimate moment, though deficits were milder in material properties such as toughness (**S. Fig. 3O-T**). Overall, the Ptgs2-Y385F mutation negatively impacted cortical bone geometry with aging to reduce bone fracture resistance.

### Ptgs2-Y385F mice have comparable load-induced bone formation and osteocyte dendritic networks to WT littermates

We quantified bone formation in response to six bouts of uniaxial forelimb compression over a period of 15 days in Ptgs2-Y385F, heterozygous, and WT mice of both sexes using dynamic histomorphometry (**Fig. 1A-C**). Relative measures of periosteal or endosteal mineralizing surface per bone surface (MS/BS), mineral apposition rate (MAR), and bone formation rate per bone surface (BFR/BS) were not significantly different between genotypes (**Fig. 2F-K**). When loaded and non-loaded limbs were considered separately, minor differences were observed. On the periosteal surface of loaded limbs, Ptgs2-Y385F mice have a slightly decreased load response driven by a decreased periosteal MS/BS in the loaded limbs as compared to WT, suggesting a possible difference in osteoblast activation on the periosteal surface (**S. Fig. 4F-H and 4L-N**). These decreases appear to be more pronounced in female Ptgs2-Y385F mice.

On the endosteal surface, loaded limbs between genotypes were not significantly different, but non-loaded limbs showed that Ptgs2-Y385F mice have significantly increased BFR/BS driven by increased MAR (**S. Fig. 4I-K and 4O-Q**). Interestingly, when sexes were analyzed separately, female mice did not show any differences due to genotype in non-loaded limbs on either bone surface, but males showed a significant decrease in non-loaded periosteal MS/BS and significant increases in non-loaded endosteal MAR and BFR/BS. Additionally, we used phalloidin staining for F-actin to observe any genetic differences in osteocyte dendritic density at 20 weeks of age. Osteocytes in cortical bone near the endosteal and periosteal surfaces were imaged and analyzed separately (**Fig. 2D-E**). Both analysis methods revealed that there were no significant differences in dendrite number per osteocyte between genotypes and only mild, non-significant, decreases in dendritic density between genotypes (**Fig. 2L-N**). No additional differences were observed in dendrite number or density when sexes were analyzed separately. Taken together, these findings demonstrate that mice carrying the Ptgs2-Y385F mutation have normal load-induced bone formation and osteocyte dendritic networks as compared to WT littermates.

### The Ptgs2-Y385F mutation does not cause significant changes to tibial morphology, osteoblast and osteocyte number, MMP-13 expression, or collagen fibril thickness

Non-loaded tibias from 16–18-week Ptgs2-Y385F and WT mice of both sexes were analyzed histologically. First, trabecular and cortical bone was stained with H&E (**Fig. 3A-D**). There were no significant differences due to genotype in growth plate thickness (**Fig. 3K**), osteoblast number (**Fig. 3L**), total lacunae number (**Fig. 3M**), or empty lacunae number between genotypes, but there was a trending decrease in empty and partially filled lacunae in Ptgs2-Y385F mice compared to WT (**Fig. 3N**). Second, MMP-13 in cortical osteocytes and growth plate chondrocytes was assessed using immunohistochemistry (**Fig. 3E-H**). There were no significant differences in MMP-13 staining in either bone compartment between genotypes (**Fig. 3O-P**). Third, tibial collagen fibril thickness was analyzed using picrosirius red staining to distinguish between thin (green), intermediate (yellow), and thick (red) fibrils in whole tibial cross-sections (**Fig. 3I-J**). Collagen fibril thickness trended lower in Ptgs2-Y385F mice compared to WT (p=0.066 by two-way ANOVA of the interaction between fibril thickness and genotype), suggesting a possible effect of the mutation (**Fig. 3Q**). No sex-based differences due to genotype were identified from histological analysis. Representative images from male mice and MMP-13 immunofluorescence controls are shown in **Supplementary Figure 5**.

**Figure 3:**
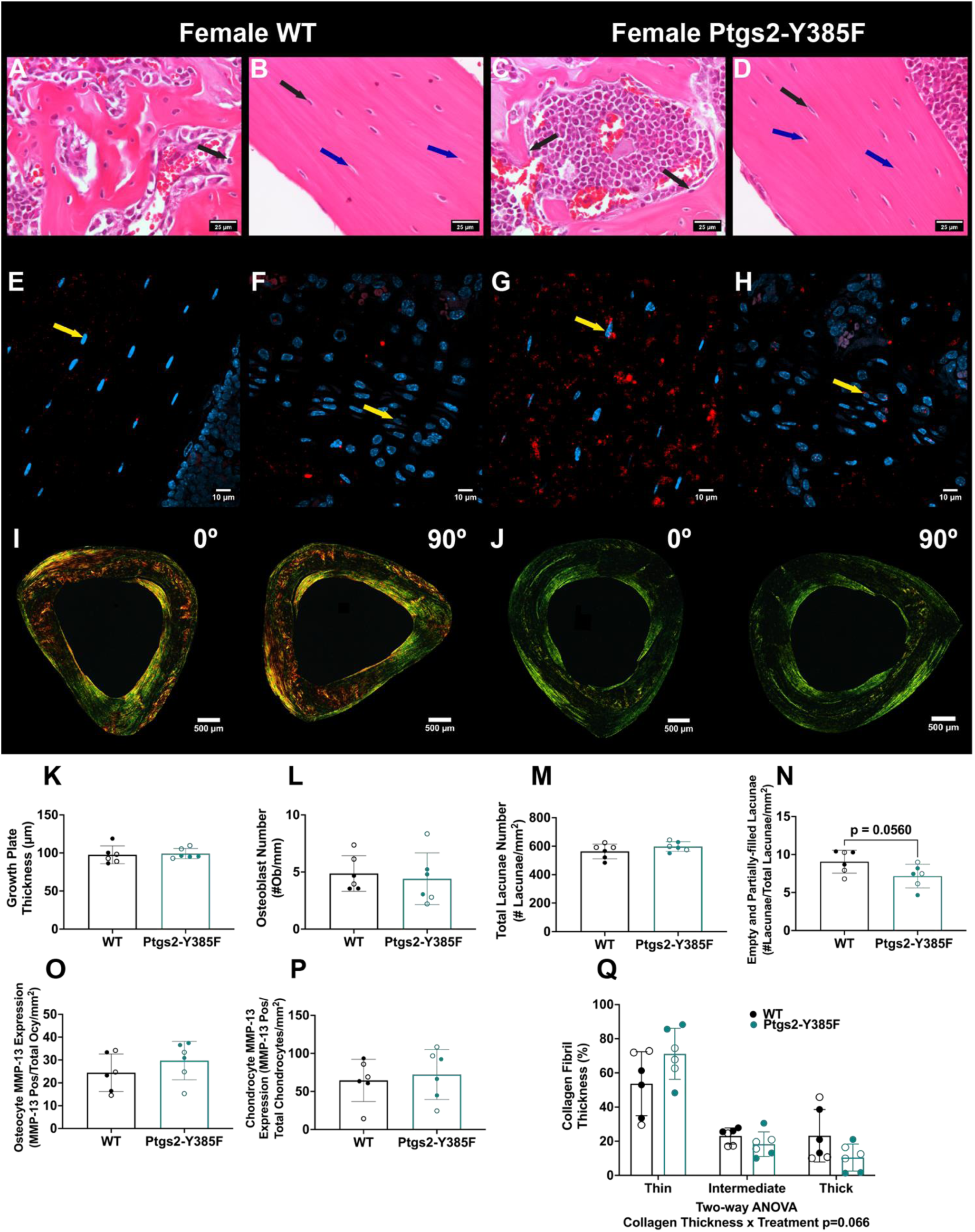
Ptgs2-Y385F mice have comparable tibial growth plate thickness, osteoblast number, osteocyte lacunae number, MMP13 expression, and collagen organization to WT littermates. (A-D) Representative longitudinal tibial sections stained with hematoxylin and eosin (H&E) of trabecular and cortical bone, (E-H) representative images of longitudinal tibial sections immunofluorescence staining for MMP-13 (red) and DAPI staining of nuclei (blue) of cortical bone and growth plate chondrocytes, and (I-J) representative images of tibial cross sections stained with picrosirius red and visualized using polarized light all from female WT and Ptgs2-Y385F mice aged 16-18 weeks at experiment end. For immunofluorescence staining, any red puncta colocalizing with DAPI staining was considered a cell expressing MMP-13. (K) Quantification of tibial growth plate thickness (representative images included in Supplemental Figure 5), (L) osteoblast number per bone perimeter in trabecular bone, (M) total osteocyte lacunae number in cortical bone, (N) number of empty or partially-filled osteocyte lacunae in cortical bone, (O) osteocyte MMP-13 expression in cortical bone, (P) chondrocyte MMP-13 expression in the growth plate, and (Q) collagen fibril thickness percentage of total stained area. n=5-6 per group. Black arrows indicate osteoblasts in trabecular bone or filled lacunae/osteocytes in cortical bone, blue arrows indicate empty or partially-filled lacunae in cortical bone, and yellow arrows indicate cells expressing MMP-13. A p value below 0.05 was considered significant and a p value below 0.1 was considered trending. Welch’s t test F statistic for H&E and MMP-13 quantifications ranged from 1.046 to 2.995. Welch’s ANOVA F statistic for picrosirius red quantifications ranged from 0.006935 to 32.42. Female replicates are displayed as filled circles and male replicates as empty circles.

### Ptgs2-Y385F mice form and repair fatigue fractures similarly to WT littermates

We subjected Ptgs2-Y385F mice and WT littermates of both sexes to fatigue loading to induce a non-displaced fracture at the ulna midpoint (**Fig. 4A-D**). We observed no significant differences in cycle number to fatigue fracture formation, fracture location, or fracture crack length due to genotype or sex (**Fig. 4E-G**). Analysis of the bone callus by microCT 15 days post-injury revealed no significant differences in woven bone mineral density, woven bone extent, or woven bone volume (**Fig. 4H-J**). Next, mechanical testing of the injured limb revealed no significant differences in ultimate load, ultimate energy, or stiffness when relative to the non-injured, contralateral limb (**Fig. 4K-M**) or when comparing limbs separately (**S. Fig. 6A-F**). We did observe small, but significant, decreases in ulnar tissue mineral density and bone area/tissue area in Ptgs2-Y385F mice compared to WT littermates in the non-injured limb (**S. Fig. 6G-I**). Ptgs2-Y385F mice showed significantly reduced tissue mineral density in both radius regions, with no genotype differences in cross-sectional geometry (**S. Fig. 6J-O**).

**Figure 4:**
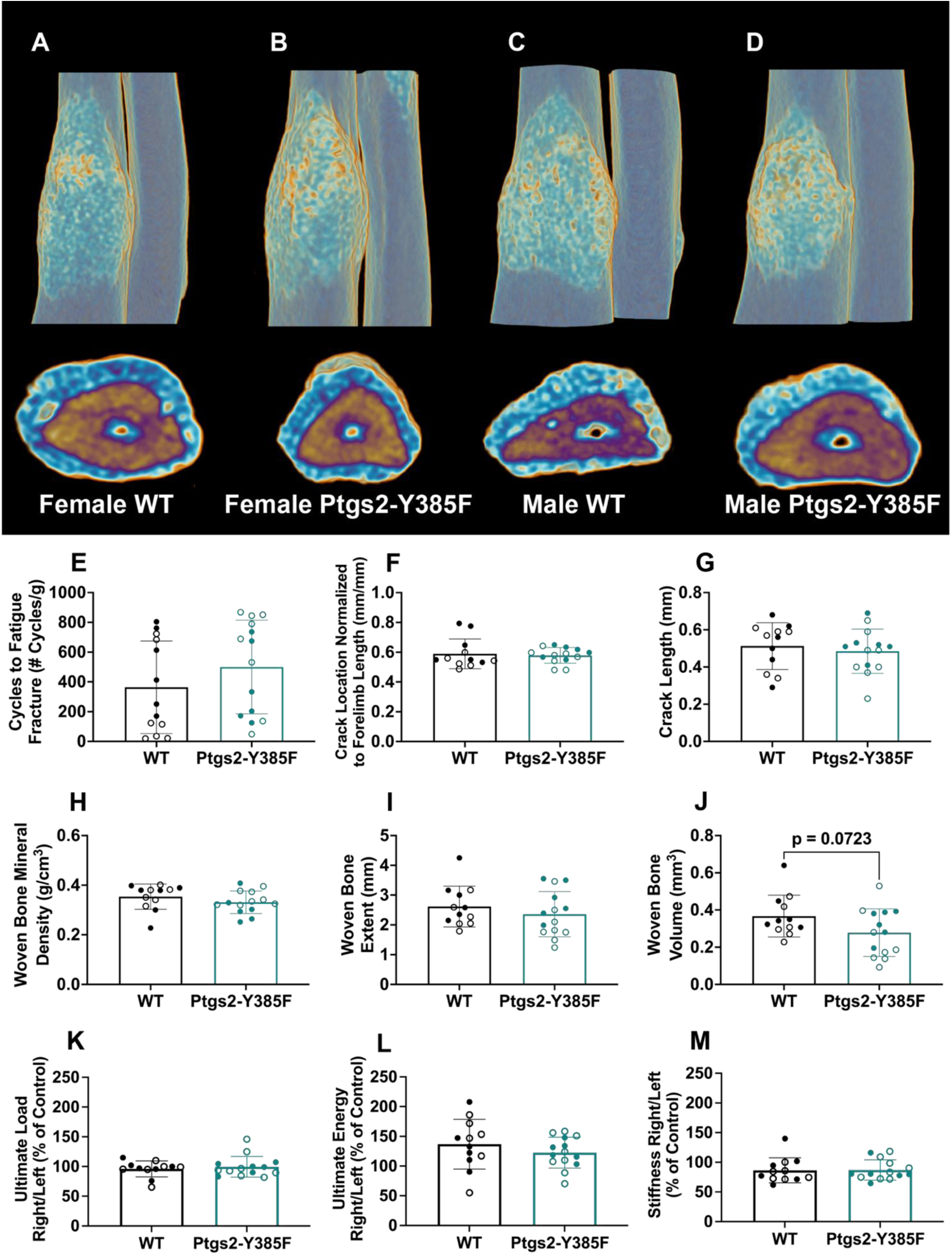
Ptgs2-Y385F mice form and heal fatigue fractures similarly to WT littermates. Fatigue fracture formation was measured using the number of cycles required to reach a calculated displacement limit, fracture healing was measured using microCT scanning of the injured limb 15 days post-fracture (experimental day 30), and the mechanical performance of the bone callus was measured using monotonic compression to failure. (A-D) Representative microCT reconstructions of female mouse forelimbs showing fracture calluses and cross sections of the injuries approximately where the fracture crack begins. (E) Quantification of the number of cycles required to reach the calculated displacement limit per mouse normalized to body mass at injury (experimental day 15). (F) Measurement of fracture crack location normalized to forelimb length, (G) measurement of fracture crack length, (H) quantification of woven bone mineral density of the callus, (I) measurement of woven bone extent along forelimb, and (J) quantification of woven bone volume. n=12-14 per group. (K) Calculated ultimate load, (L) ultimate energy, and (M) stiffness of the injured limb (right) relative to the intact limb (left). n=12-14 per group. A p value below 0.05 was considered significant and a p value below 0.1 was considered trending. Welch’s t test F statistic for these analyses ranged from 1.019 to 3.591. Female replicates are displayed as filled circles and male replicates as empty circles.

### Naproxen reduces load-induced bone formation in WT but not Ptgs2-Y385F mice and decreases bone toughness in both genotypes

We next used the Ptgs2-Y385F model to determine the COX2-dependent and -independent effects of naproxen. Ptgs2-Y385F and WT mice received naproxen sodium (10.9 mg/kg) or vehicle drinking water for 15 days. During this, each mouse underwent 6 bouts of non-damaging axial forelimb compression. First, load-induced bone formation was quantified by dynamic histomorphometry (**Fig. 5A-D, S. Fig 7A-D**). WT mice displayed a decrease in relative periosteal bone formation rate per bone surface (rPs.BFR/BS) with treatment, but Ptgs2-Y385F mice did not (**Fig. 5E-G**). Analysis of loaded and non-loaded forelimbs separately showed that naproxen decreased MS/BS, MAR, and BFR/BS in the loaded limb on the periosteal surface in only WT, but not Ptgs2-Y385F mice (**S. Fig. 7E-G**). However, there were no significant differences due to treatment on the endosteal surface (**Fig. 5H**) or in non-loaded limbs (**S. Fig. 7**). Further, naproxen did not significantly decrease measures of trabecular or cortical geometry (**Fig. 5I-K**), although some parameters had small, but significant, increases due to treatment after 15 days. However, there were no significant differences detected after treating for 30 days (**S. Fig. 8, 9**). Crucially, analysis by standard three-point bending demonstrated that naproxen treatment for 15 days significantly decreased ultimate strain, post-yield energy, ultimate bending energy, and toughness in both genotypes without decreasing ultimate moment or Young’s modulus (**Fig. 5L-Q**). Surprisingly, 30 days of naproxen treatment impaired mechanical performance in female WT mice, with milder effects in Ptgs2-Y385F females. In contrast, WT males showed no change or slight improvements, while Ptgs2-Y385F males had mild decreases (**S. Fig. 8G-P**).

**Figure 5:**
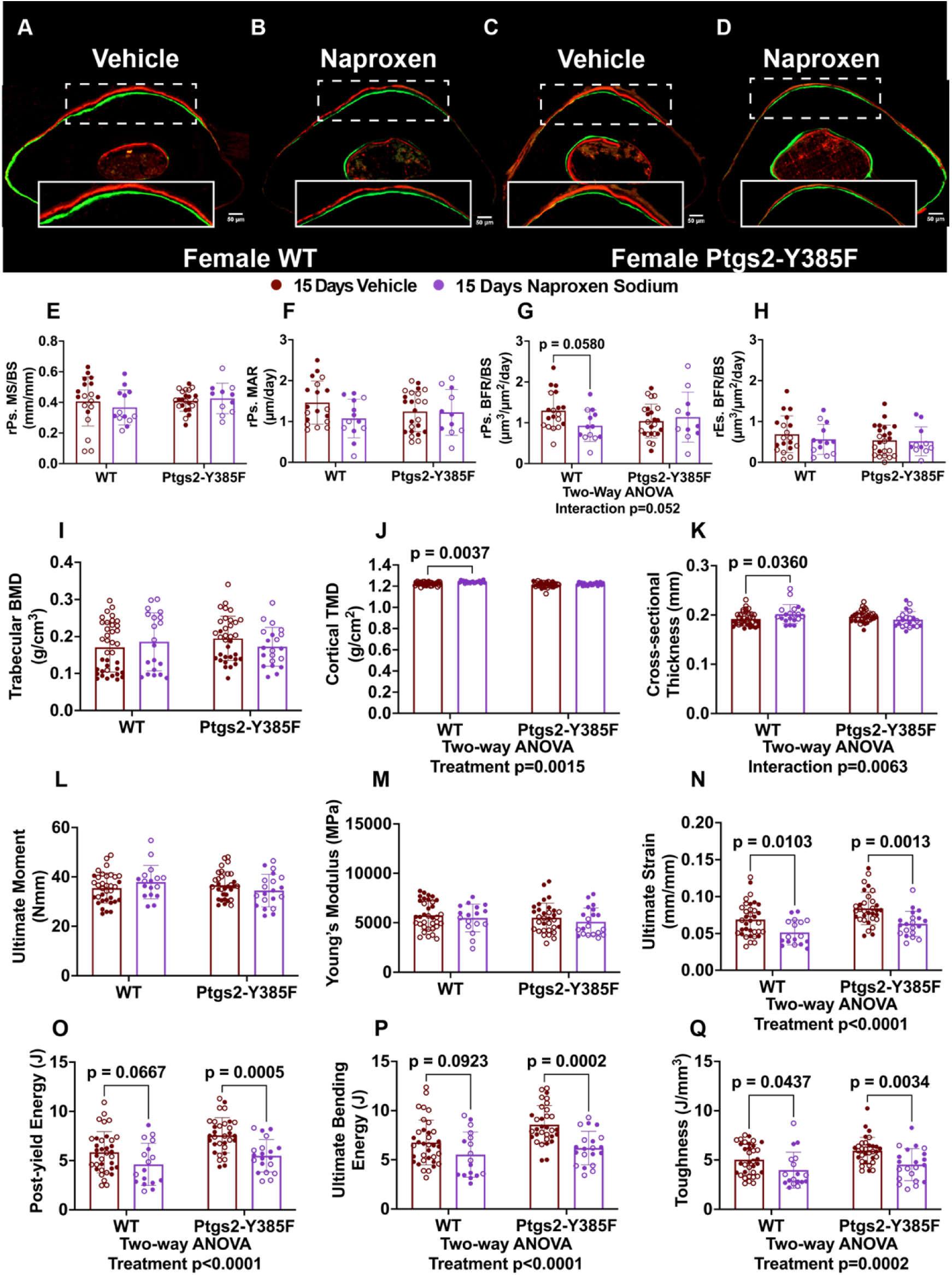
Naproxen treatment for 15 days decreases load-induced bone formation in only WT mice, but decreases bone toughness in both genotypes without affecting bone geometry or mineral content. Bone formation in response to uniaxial forelimb compression was measured in ulnar mid-diaphysis sections using dynamic histomorphometry with calcein (green) and alizarin red (red) bone formation labels in mice aged 16-18 weeks at experiment end who received vehicle or naproxen drinking water for 15 days. Bone geometry was quantified using microCT scanning and reconstruction of non-loaded femurs of the same mice and femoral mechanical performance was measured using standard three-point bending. (A-D) Representative sections from female WT and Ptgs2-Y385F mice that received either vehicle or naproxen drinking water. Dynamic histomorphometry outcomes included (E) relative periosteal mineralizing surface per bone surface (rPs. MS/BS), (F) relative periosteal mineral apposition rate (rPs. MAR), (G) relative periosteal bone formation rate per bone surface (rPs. BFR/BS), and (H) relative endosteal bone formation rate per bone surface (rEs BFR/BS). n=11-24 per group. Bone geometry parameters include (I) trabecular bone mineral density (BMD), (J) cortical tissue mineral density (TMD), and (K) cortical cross-sectional thickness. n=20-38 per group. Mechanical performance parameters include (L) ultimate moment, (M) Young’s modulus, (N) ultimate strain, (O) post-yield energy, (P) ultimate bending energy, and (Q) toughness. n=17-35 per group. A p value below 0.05 was considered significant and a p value below 0.1 was considered trending. Welch’s ANOVA F statistic for dynamic histomorphometry measures ranged from 0.04185 to 3.938. Welch’s ANOVA F statistic for microCT and standard 3-point bending measurements ranged from 0.009907 to 18.01. Female replicates are displayed as filled circles and male replicates as empty circles

Finally, a separate cohort of female C57BL/6J mice were given the same “low dose” of naproxen (10.9 mg/kg/day), or a “high dose” naproxen (41.6 mg/kg/day) for 30 days. Bone geometry analysis by microCT showed significant, but modest, decreases in trabecular thickness and cortical bone area/tissue area with high dose naproxen compared to low dose naproxen. Importantly, there were no significant differences between doses in ultimate moment, Young’s modulus, or toughness (**S. Fig. 8Q-X**).

### Naproxen alters osteoblast number, osteocyte number and viability, and measures of osteocyte perilacunar canalicular remodeling (PLR)

Non-loaded tibias from 16–18-week Ptgs2-Y385F and WT mice of both sexes given vehicle or naproxen (10.9 mg/kg) for 15 days were analyzed histologically, first using H&E (**Fig. 6A-D**). Osteoblast number per bone perimeter in trabecular bone was significantly increased in WT, but not Ptgs2-Y385F, mice given naproxen compared to vehicle (**Fig. 6G**). In cortical bone, total osteocyte lacunae number was significantly increased in only WT mice with treatment; empty lacunae number, a measure of osteocyte death, was significantly increased in both genotypes due to treatment (**Fig. 6H-I**). We note that the effects of naproxen on osteocytes appear more pronounced in female than male mice. No significant differences due to treatment were observed in tibial growth plate thickness, but naproxen significantly increased the number of empty and partially filled osteocyte lacunae in Ptgs2-Y385F, but not WT, mice (**S. Fig. 10K-L**). Next, MMP-13 was analyzed using immunohistochemistry in cortical osteocytes from female WT and Ptgs2-Y385F mice (**Fig. 6A-D**). There were no significant differences due to treatment in the number of cortical osteocytes positive for MMP-13 (**Fig. 6J**), but there was an increase in growth plate chondrocytes positive for MMP-13 in WT, but not Ptgs2-Y385F, mice (**S. Fig. 10M**). Finally, fluorophore-labeled osteocytes from female WT mice given vehicle or naproxen were analyzed (**Fig. 6E-F**). There was a significant increase in the number of fluorophore-labeled osteocytes near the endosteal, but not the periosteal, surface of ulnar bone (**Fig. 6J-K**). These sections showed a trend of decreased empty osteocyte lacunae near the endosteal surface, but increased number of empty osteocyte lacunae near the periosteal surface (**S. Fig. 10N-O**).

**Figure 6:**
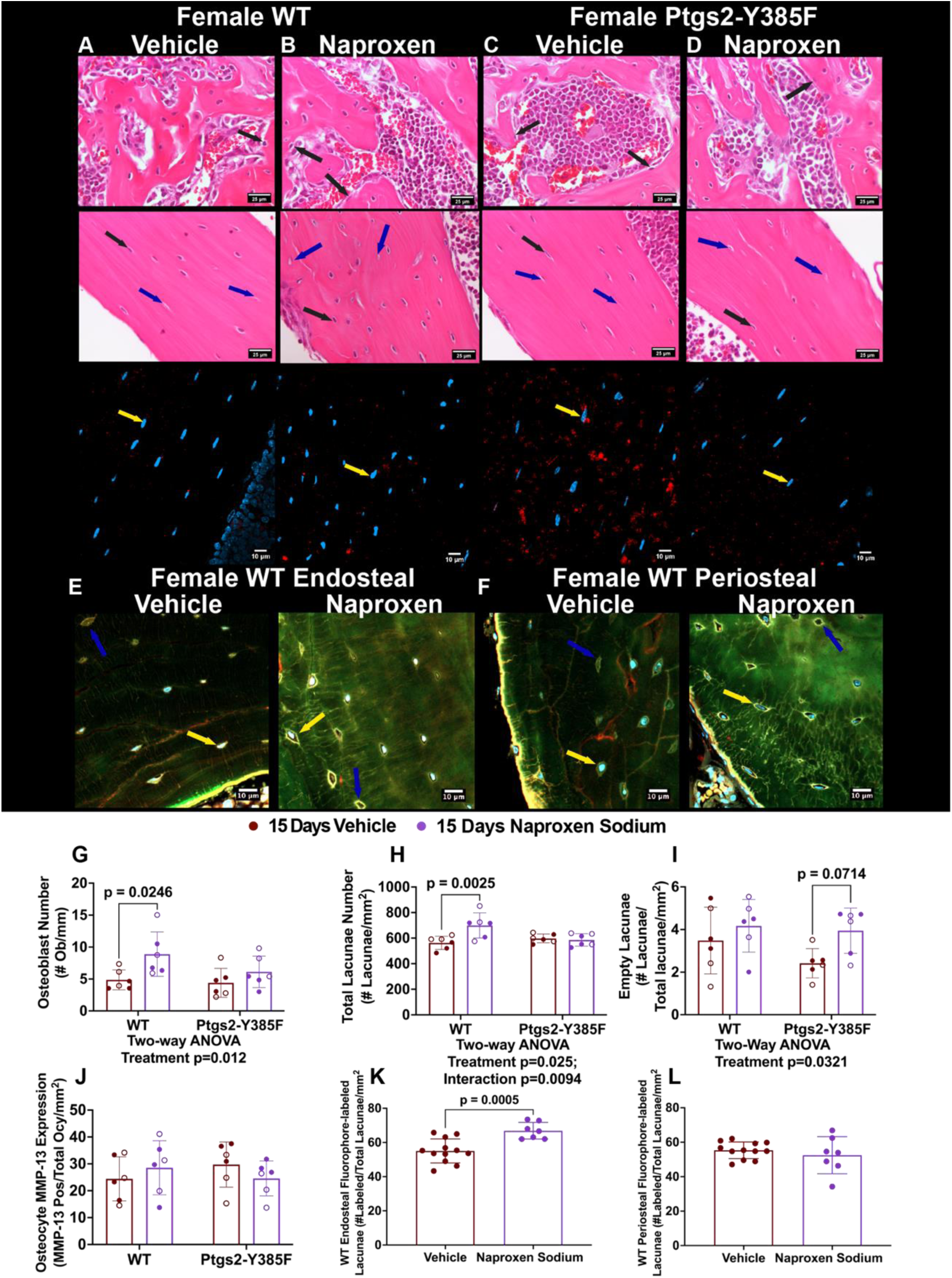
15 days of naproxen treatment increases osteoblast number and osteocyte lacunae number in only WT mice, increases empty lacunae number in both genotypes, and may increase osteocyte PLR in WT mice. (A-D) Representative longitudinal tibial sections stained with hematoxylin and eosin (H&E) of trabecular and cortical bone and representative images of longitudinal tibial sections immunofluorescence staining for MMP-13 (red) and DAPI staining of nuclei (blue) of cortical bone osteocytes from female WT and Ptgs2-Y385F mice aged 16-18 weeks at experiment end that received vehicle or naproxen drinking water for 15 days. For immunofluorescence staining, any red puncta colocalizing with DAPI staining was considered a cell expressing MMP-13. (E-F) Representative sections of forelimbs used for dynamic histomorphometry and stained with DAPI to visualize osteocyte fluorophore labeling in cells near the endosteal and periosteal surfaces of bone from female WT mice that received vehicle or naproxen drinking water for 15 days. (G) Quantification of osteoblast number per bone perimeter in trabecular bone, (H) total osteocyte lacunae number in cortical bone, (I) number of empty osteocyte lacunae in cortical bone, (J) osteocyte MMP-13 expression in cortical bone. n=5-6 per group. Quantification of osteocytes labeled with either calcein, alizarin red, or both labels near the (K) endosteal and (L) periosteal surfaces of the bone. n=7-12 per group. Black arrows indicate osteoblasts in trabecular bone or filled lacunae/osteocytes in cortical bone, blue arrows indicate empty or partially-filled lacunae in cortical bone, and yellow arrows indicate cells expressing MMP-13 in immunofluorescence staining or labeled with fluorophores in sections from dynamic histomorphometry. A p value below 0.05 was considered significant and a p value below 0.1 was considered trending. Welch’s ANOVA F statistic for H&E and MMP-13 quantifications ranged from 0.0235 to 8.247. Welch’s t test F statistic for fluorophore labeled osteocyte quantifications ranged from 2.083 to 4.962. Female replicates are displayed as filled circles and male replicates as empty circles. Black arrows indicate osteoblasts in trabecular bone or full osteocyte lacunae in cortical bone, blue arrows indicate empty or partially-filled osteocyte lacunae in cortical bone, yellow arrows indicate osteocytes staining positive for MMP-13 immunofluorescence or fluorophore-labeled osteocyte lacunae containing a nucleus.

### Naproxen treatment changes collagen fibril thickness in Ptgs2-Y385F mice and alters osteocyte dendritic networks in both genotypes

Collagen fibrils were visualized with polarized light images of mid-diaphyseal, non-loaded tibia sections stained with picrosirius red from mice treated with vehicle or naproxen (10.9 mg/kg) for 15 days (**Fig. 7A-D**). Quantifications of thin (green), intermediate (yellow), and thick (red) collagen fibrils revealed that naproxen decreased thin collagen fibrils and increased intermediate and thick fibrils in Ptgs2-Y385F mice compared to vehicle controls (**Fig. 7E-G**). Analyzing collagen fibril organization with sexes separate revealed no sex-dependent differences. Brightfield microscopy images of picrosirius red staining for all treatment groups and sexes were consistent with these findings (**S. Fig. 12**). Mid-diaphyseal cross sections from non-loaded tibia of WT mice receiving vehicle or naproxen (10.9 mg/kg) for 30 days were similarly analyzed. WT mice of either sex did not show significant differences in collagen fibril thickness with treatment, though there was a trend of treatment to diminish thick fibrils (**S. Fig. 11E-G**).

**Figure 7:**
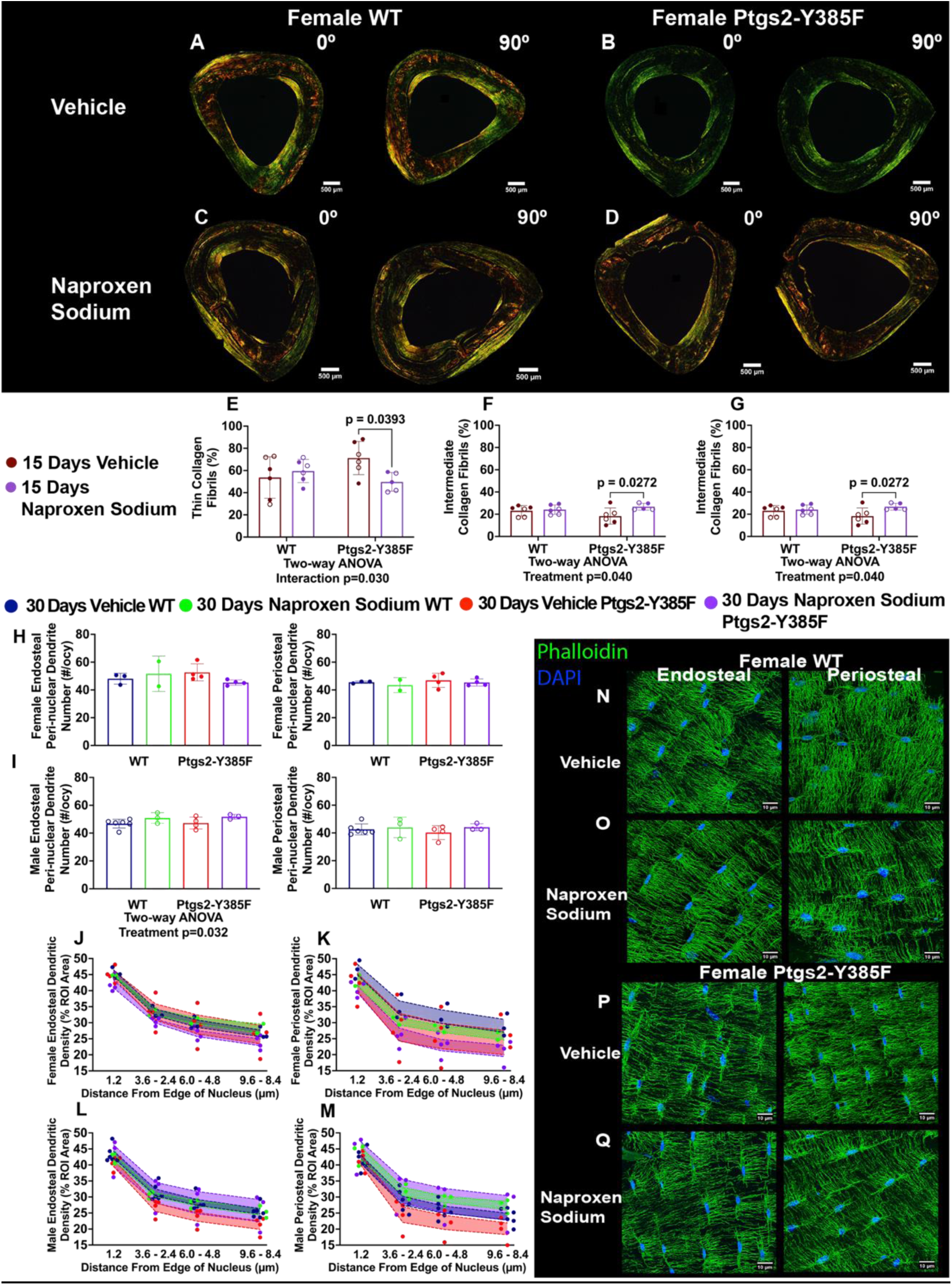
Naproxen treatment alters collagen thickness in Ptgs2-Y385F mice and has sexually dimorphic effects on osteocyte dendritic networks. Changes to collagen fibril thickness with treatment were quantified using picrosirius red stained cross sections of tibias from mice treated with vehicle or naproxen for 15 days and imaged using polarized light. Osteocyte dendritic networks were visualized in mid-cortical sections of non-loaded femurs using phalloidin F-actin staining (green) and DAPI nuclear staining (blue), and networks were quantified using dendrite counting near the nucleus (peri-nuclear) and at a standard distance away from the nucleus where dendrites commonly ended (terminal) and dendritic density was measured using the moving band method in mice aged 20 weeks at experiment end that received vehicle or naproxen treatment for 30 days. (A-D) Representative images of tibial cross sections stained with picrosirius red from female WT and Ptgs2-Y385F mice aged 16-18 weeks at experiment end that received vehicle or naproxen for 15 days. Quantifications of (E) thin (green), (F) intermediate (yellow), and (G) thick (red) collagen fibrils measured as the percentage of stained area in the cross section. n=5-6 per group. Dendrite number per osteocyte in the peri-nuclear region for (H) female and (I) male WT and Ptgs2-Y385F mice. Dendritic density as measured by the moving band method in for female osteocytes near the (J) endosteal and (K) periosteal surfaces of the bone and (L-M) the corresponding measurements in osteocytes from male mice. The boundaries of the standard error of the mean are colorized alongside individual replicates. n=2-4 for female mice and n=3-6 for male mice. (N-Q) Representative sections from female WT and Ptgs2-Y385F mice that received vehicle or naproxen treatment for 30 days showing the dendritic network near the endosteal and the periosteal surfaces of the bone. A p value below 0.05 was considered significant and a p value below 0.1 was considered trending. Welch’s ANOVA F statistic for these quantifications ranged from 0.01613 to 5.474. Female replicates are displayed as filled circles and male replicates as empty circles.

Additionally, osteocyte dendritic networks were visualized in mid-diaphyseal cross sections from non-loaded femurs of mice treated for 30 days (**Fig. 7N-Q**). Dendrite number per osteocyte in the peri-nuclear region, representing dendrite sprouting from the cell body, was increased in both male WT and Ptgs2-Y385F mice with naproxen, but no changes were observed in female mice in osteocytes near the endosteal surface of bone (**Fig. 7H-I**). Dendritic density measured by the “moving band method” showed more pronounced effects of naproxen on dendritic density in osteocytes near the periosteal surface compared to the endosteal surface in both sexes, and demonstrated opposing trends in dendritic density between sexes (**Fig. 7J-M**). Representative images of male samples are shown in **Supplementary Figure 11**.

### Naproxen pre-treatment has sexually dimorphic effects on fatigue fracture formation and continued treatment during fracture healing decreases bone callus volume in female, but not male, mice of both genotypes

Mice of both genotypes and sexes were pre-treated with vehicle or naproxen (10.9 mg/kg) for 15 days before undergoing fatigue injury by cyclical loading. Longitudinal and cross-sectional reconstructions of microCT imaging were used to analyze the effect of naproxen on fracture calluses from WT and Ptgs2-Y385F mice, both female (**Fig. 8A-D**) and male (**S. Fig. 13A-D**). Female WT mice given naproxen showed a decreased cycle number to fracture compared to vehicle, whereas Ptgs2-Y385F females showed an increase (**Fig. 8E**). Contrastingly, Male WT mice showed moderate increases in cycle number with treatment whereas Ptgs2-Y385F mice showed decreases in cycle number (**Fig. 8F**). Calculations of instantaneous stiffness showed corroborating decreased stiffness in WT females and increased stiffness in Ptgs2-Y385F females with naproxen, but showed no significant differences due to treatment in males (**Fig. 8G-H**).

**Figure 8:**
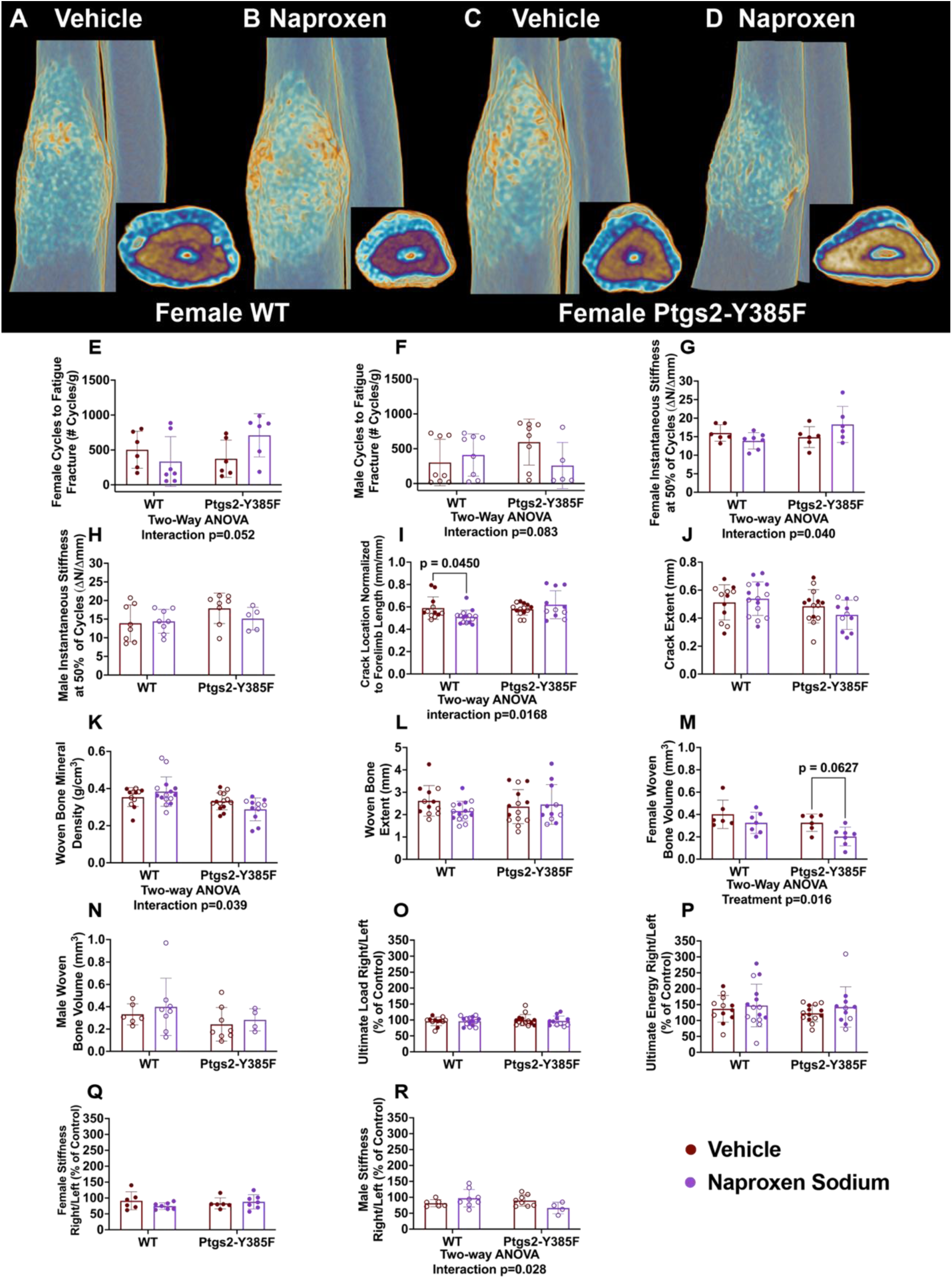
Naproxen treatment has sexually dimorphic effects on fatigue fracture formation and healing in both WT and Ptgs2-Y385F mice. Fatigue fracture formation was measured using the number of cycles required to reach a calculated displacement limit significant fatigue fracture formation, forelimb instantaneous stiffness during fracture formation was measured using the difference in measured force divided by the difference in measured displacement during compression, fracture healing was measured using microCT scanning of the injured limb 15 days post-fracture (experimental day 30), and the mechanical integrity of the bone callus was measured using monotonic compression to failure. (A-D) Representative microCT reconstructions of female mouse forelimbs after 30 days of treatment showing fracture calluses and cross sections of the injuries approximately where the fracture crack begins. Quantification of the number of cycles required to reach the calculated displacement limit per mouse normalized to body mass at injury (experimental day 15) for (E) female and (F) male mice. Instantaneous stiffness calculated at 50% of total cycles per mouse for (G) female and (H) male mice. (I) Measurement of fracture crack location normalized to forelimb length, (J) measurement of fracture crack length, (K) quantification of woven bone mineral density of the callus, (L) measurement of woven bone extent along forelimb, and quantification of woven bone volume for (M) female and (N) male mice. n=12-14 per group. (O) Calculated ultimate load, (P) ultimate energy, and stiffness of (Q) female and (R) male mice of the injured limb (right) relative to the intact limb (left). n=12-14 per group. A p value below 0.05 was considered significant and a p value below 0.1 was considered trending. Welch’s ANOVA F statistic for these measurements ranged from 0.01121 to 11.78. Female replicates are displayed as filled circles and male replicates as empty circles.

Forelimbs were excised 15 days post injury and scanned using microCT to quantify woven bone geometry and callus size. Fracture crack location was significantly altered in WT, but not Ptgs2-Y385F, mice treated with naproxen as compared to vehicle, indicating that injuries were more likely to form proximally with naproxen treatment (**Fig. 8I**). There was no significant difference in fracture crack extent across treatment groups and genotypes (**Fig. 8J**). No significant differences were observed due to treatment in woven bone mineral density or woven bone extent, but there was a significant decrease in woven bone volume across genotypes in female mice with naproxen compared to vehicle, indicating sexual dimorphism in fatigue fracture healing with treatment (**Fig. 8K-N**). The mechanical performance of these forelimbs was measured using monotonic compression to failure. Ultimate load and ultimate energy of the injured limb relative to the intact limb were not altered, but there was an increase in male WT relative stiffness and a decrease in male Ptgs2-Y385F relative stiffness with naproxen that was not seen in females (**Fig. 8O-R**). Mechanical performance of the injured and intact limbs was examined separately. Here, intact limbs showed no significant differences due to treatment in any measure across sexes and genotypes besides increased ultimate energy in only male WT mice with naproxen treatment compared to controls (**S. Fig. 13E-J**). Injured limbs showed some significant differences due to treatment, mainly in female WT mice (**S. Fig. 13K-P**).

## DISCUSSION

We determined COX2-independent and COX2-dependent effects of the non-selective NSAID naproxen in bone under basal, non-damaging loading, and damaging loading conditions using the Ptgs2-Y385F mouse model. We hypothesized that naproxen would diminish strain adaptive bone remodeling in a COX2-dependent manner while deleteriously affecting osteocytes and bone toughness in a COX2-independent manner. Consistent with this hypothesis, our results demonstrate that naproxen decreased strain adaptive bone remodeling in only WT mice and decreased bone toughness in both WT and Ptgs2-Y385F mice of both sexes. Therefore, we conclude that diminished bone toughness from naproxen treatment is a COX2-independent effect. However, the effects of naproxen on osteocyte viability, dendritic networks, and remodeling activity are evidently a combination of COX2-dependent and COX2-independent mechanisms. Further, fatigue fracture experiments did not demonstrate a large effect size of naproxen pretreatment on fatigue fracture induction as would be expected based on femoral bone toughness measurements. Significant results from these experiments support a more detrimental effect of naproxen on fatigue fracture induction and cortical bone healing in females than males. Additionally results suggest that the mechanisms underpinning the published correlations between regular NSAID use and stress fracture are likely not limited to NSAID inhibition of COX2, as Ptgs2-Y385F mice also displayed differences due to treatment. Altogether, our findings suggest that regular use of a COX2-selective NSAID for pain relief may not be sufficient to minimize fracture risk as NSAID effects in bone have both COX2-dependent and COX2-independent mechanisms.

We first validated the Ptgs2-Y385F mouse model as a usable comparison to WT littermates at skeletal maturity. Results demonstrated that this model does not display the same stunted skeletal growth as the COX2^-/-^ mouse, and did not require strain matching or adjustments for body mass to compare to WT mice during mechanical loading^14,60,61^. The mutation also did not decrease femoral bone mechanical performance at either the 16-18 week or 20 week timepoints, unlike the COX2^-/-^ mouse^61^. Our results aligned with work by Robertson et al. showing mild differences in bone geometry and increased work to fracture of male, but not female, COX2^-/-^ mice, indicating that mouse genetic backgrounds may influence bone development with genetic COX2 inhibition^62^. Analyses of bone geometry and mechanical performance were repeated in an aged cohort of male Ptgs2-Y385F and WT mice from our colony. WT males showed expected changes in trabecular and cortical bone with aging^63,64^. Surprisingly, Ptgs2-Y385F males displayed stunted cortical bone geometry adaptation with aging, suggesting a difference in the modeling processes of endosteal and periosteal surfaces in this model. This observation should be explored further as elderly individuals are more likely to be regularly using NSAIDs and changes in bone modeling could contribute to increased fracture risk^65^.

Second, we subjected WT, heterozygous, and Ptgs2-Y385F mice of both sexes to non-damaging uniaxial forelimb compression. No significant differences in relative measures of load-induced bone formation due to genotype were found, but this model likely has differences in mechanosensation or transduction mechanisms that were evident when loaded and non-loaded limbs were analyzed separately. These results agree with previous literature establishing that a functional COX2 enzyme is not required for lamellar bone formation^61^. The Ptgs2-Y385F mouse likely compensates for COX2 deficiency with other mechanosensory mechanisms or through heterodimer formation of the mutated COX2 enzyme and intact COX1 enzyme to produce similar prostaglandin signaling^14,50,61,66–74^. To investigate this orthogonally, we analyzed the morphology of the osteocyte dendritic network as it is a major mechanosensor in bone^47,63,66,67,75–78^. We saw no differences in the number of dendrites per osteocyte, and only mild decreases in dendritic density due to genotype. This suggests that changes to the dendritic network caused by the mutation are not solely responsible for differences in mechanosensation.

Third, we analyzed non-loaded tibias from WT and Ptgs2-Y385F mice of both sexes histologically. We saw no significant differences between WT and Ptgs2-Y385F mice in any measure. We did see an almost significant decrease in the number of empty (dead osteocytes) and partially-filled (dying osteocytes) lacunae in cortical bone of Ptgs2-Y385F mice compared to WT controls, suggesting a potential decrease in osteocyte apoptosis in Ptgs2-Y385F mice. This finding should be confirmed with direct measures of apoptosis. We also observed non-significant alterations in collagen fibril thickness through picrosirius red staining due to genotype that should be confirmed with further collagen analysis methods. This adjustment in collagen with COX2 inhibition agrees with our previously observed changes in picrosirius red staining with naproxen treatment in C57BL/6J females, but as this same change was not seen in the WT or Ptgs2-Y385F mice with naproxen treatment, it is unclear if the same mechanism is responsible^7^.

Fourth, a separate cohort of WT and Ptgs2-Y385F mice received fatigue fracture loading, and we analyzed fracture formation, healing, and post-healing mechanical performance. There were no significant differences due to genotype on stress fracture formation or post-healing mechanical performance of the forelimb, but a non-significant decrease in woven bone volume of the fracture callus in Ptgs2-Y385F mice compared to WT controls was observed. This finding aligns with the known importance of COX2-produced prostaglandins for fracture healing. Its non-significant effect size suggests that Ptgs2-Y385F mice have compensatory mechanisms for their COX2 deficiency during healing that should be explored further by probing the contents of the fracture callus^7,25^.

The analyses described above were repeated with mice receiving vehicle or naproxen treatment for 15 days in non-damaging loading or 30 days in damaging loading experiments. To increase the clinical relevance of this study, a naproxen dose of 10.9 mg/kg/day, translating to a human dose of about 53 mg for a 60kg adult, was used for most studies and when a high dose was used (41.6 mg/kg/day), it was within the daily recommended dose, translating to about 203 mg for a 60 kg adult. As naproxen is not commonly sold below 220 mg, both doses used are realistic and conservative.

Critically, we found that naproxen treatment for 15 days decreased lamellar bone formation in WT, but not Ptgs2-Y385F, mice but decreased femoral mechanical performance in both genotypes. This decrease in mechanical performance was observed without significant decreases in cortical bone geometry measures or young’s modulus. Thus, we conclude that naproxen’s effects on lamellar bone formation are through its inhibition of COX2 and the production of PGE2, but that its effects on bone toughness are due to a COX2-independent mechanisms displayed by both male and female mice^24,46^. This finding adds to a previously proposed mechanism of increased stress fracture risk with NSAID use due to reduced strain adaptive bone remodeling^6^. These results also suggest a non-COX1 compensation during mechanical loading by Ptgs2-Y385F mice as their lamellar bone formation was not decreased with naproxen despite its non-selective behavior. This contradicts previous reports from the COX2^-/-^ mouse showing COX1 compensation COX2 loss during lamellar bone formation. Given that COX2^-/-^ mice die upon receiving non-selective NSAIDs and Ptgs2-Y385F mice do not, they likely have physiological differences that influence mechanosensation mechanisms^61^.

Interestingly, when a cohort receiving vehicle or naproxen for 30 days was analyzed, it was found that longer treatment with naproxen caused sexually dimorphic differences in bone. Female WT mice displayed decreases in bone area with naproxen and male WT mice showed increases with treatment, whereas Ptgs2-Y385F males and females seemed similarly unaffected by treatment. This difference was reinforced with mechanical performance measures as WT females showed decreased bone toughness with treatment and males did not. This suggests an adaptation over the treatment period in male, but not female mice, that could be at least partially linked to cortical bone geometry changes. As females are inherently more likely to form stress fractures, male, but not female, adaptation during NSAID treatment should be investigated further for potential therapeutic interventions in females^75^. Given that detrimental, but mild, effects of naproxen on bone geometry were seen in both WT females and C57BL/6J females with 30 days, but not 15 days, of naproxen treatment, it is also likely that exposure time affects bone more significantly than dose within the daily recommended dose range. This data aligns with previous literature as it is known that the risk of cardiovascular, renal, gastrointestinal, and central nervous system side-effects increases with chronic NSAID use, all of which can influence bone health and quality^76,79–85^.

We next investigated the effects of naproxen on bone histologically. Our results suggest that naproxen treatment increases osteocyte apoptosis and that WT, but not Ptgs2-Y385F, bone adjusts for this loss by differentiating more osteoblasts into osteocytes. Previous literature has shown increased osteoblast to osteocyte transition and increased intercortical fluorophore labeling in models of altered matrix metalloproteinase activity. Thus, it is possible that MMP-13 activity, or that of another MMP, could be liberating latent TGF-beta trapped in the bone matrix to shunt more osteoblasts down the osteocyte pipeline and compensate for naproxen’s apparent increase in osteocyte apoptosis^86,87^. Given that the COX2 signaling pathway is heavily involved in the regulation of MMP expression and activity, it is possible that NSAID treatment could influence extracellular matrix remodeling and cellular behavior in this way^88,89^.These data do not agree with previous *in* vitro findings. Studies examining the direct effects of NSAIDs on osteoblast differentiation show reduced differentiation with treatment, but it is possible that inter-NSAID variability and limitations inherent with *in vitro* studies can explain this disagreement^17,90,91^. Our lab previously observed changes in collagen fibril thickness with naproxen treatment, thus we repeated this analysis in WT and Ptgs2-Y385F mice treated for 15 days^7^. Unexpectedly, we saw no significant changes in collagen fibril thickness as measured by picrosirius red staining in WT mice, but did see significant decreases in thin fibrils and increases in intermediate and thick fibrils in Ptgs2-Y385F mice with treatment. This indicates a COX2-independent effect of naproxen on collagen organization. Furthermore, since collagen is a critical component of post-yielding behavior in bone, this alteration could influence fracture risk and must be investigated further using other methods to analyze collagen content, organization, and cross-linking^92^. We generally saw mild effects of naproxen on the osteocyte dendritic network, but these effects were sexually dimorphic. Our analysis would benefit from further investigation by more rigorous network analysis, but sex-dependent differences in neuron dendrite branching, thickness, and pruning under models of COX2 inhibition have been demonstrated and could manifest similarly in osteocyte dendrites^93^. The mechanism behind changes in the dendritic network is unclear, but previous models of disrupted dendritic networks show a correlation with reduced bone fracture resistance^41,94,95^.

Finally, we performed fatigue fracture studies. Naproxen pretreatment had minimal effect on fatigue fracture formation and the discrepancy between these data and those from standard 3-point bending could be due to the mechanical and geometric differences between the femur and radius/ulna that distribute strain differently^96–98^. MicroCT analysis of the fracture callus showed decreased woven bone volume in females, but not males, supporting previous findings of reduced bone healing with NSAID use^7,30^.

There are several limitations to this study. First, the COX2 deficiency of the Ptgs2-Y385F mouse is lifelong and systemic, allowing for compensatory mechanisms to develop that may confound the identification of NSAID effects as COX2-independent or –dependent. An additional comparison that would add clarity would be an inducible and reversible COX2 deficiency only during the period of NSAID administration^99^. Also, Ptgs2-Y385F mice are born at a low frequency, so to achieve robust sample sizes for comparison, these studies were conducted over several years, increasing the chance of biological variability between mice in the same cohorts^14,100^. Second, naproxen is a non-selective NSAID that strongly inhibits both COX enzymes, thus the use of a COX2-selective NSAID in conjunction with this study would clarify whether our observed COX2-independent effects were due to inhibition of COX1. Third, drinking water as a drug delivery method has known limitations, and a method more representative of human experience would deliver a calculated bolus of medicine, such as NSAID-filled mealworms^101^. To mitigate differences in drug dosing, we did monitor water intake, but future work should also analyze circulating and urinary prostaglandins to ensure equal dosing between mice. Fourth, the osteocyte dendritic network analysis methods used were high-throughput, and low cost, but lacked resolution. Additionally, our methods analyze only the morphology of F-actin and not the lacunocanalicular network. By comparing the channels in bone that likely take longer to remodel than the cytoskeleton, we would have a better understanding of the pruning or extension of dendrites in response to treatment. Fifth, the fatigue fracture model used in this study is not directly comparable in timing or mechanics to stress fracture formation in humans.

Our results open several avenues of future directions. First, identification of the mechanism by which Ptgs2-Y385F mice compensate for COX2 deficiency during mechanical loading would increase understanding of mechanosensation in bone under chronic COX2 cyclooxygenase inhibition. Second, whether NSAID use cessation reverses or mitigates alterations in bone mechanosensation and fracture resistance should be studied. Third, further investigation into how naproxen changes stress fracture healing by looking at fracture callus cellular, mineral, and organic phase content, as well as allowing fractures to fully heal and testing bone mechanical performance after healing would determine whether NSAID treatment has lasting effects on mechanical performance. Fourth, other NSAIDs should be examined to identify whether structural differences influence their effects in bone when administered at comparable, clinically relevant doses in the same mouse line. We conclude that individuals with higher fracture risk should minimize exposure to naproxen and likely also COX2-selective NSAIDs for pain relief until further investigation is done into the skeletal effects of NSAIDs.

## CONCLUSION

Regular use of naproxen affects bone through both COX2-dependent and COX2-independent mechanisms that negatively impact strain adaptive bone remodeling, bone toughness, and osteocyte viability. Collectively, the data suggest that chronic administration of NSAIDs, whether non-selective or COX2-selective, may elevate the risk of skeletal fatigue injuries. As a result, NSAID use should be minimized in individuals at high risk for fatigue-related skeletal injuries and alternative drug classes or administration methods without these effects in bone must be uncovered to address musculoskeletal pain.

## ACKNOWLEDGEMENTS

Funding for this study was provided by the Department of Orthopaedic Surgery at Thomas Jefferson University and the Rothman Orthopedic Institute. The laboratory was also supported by NIH NIAMS & NIDCR (award numbers AR052273 to AC, AR074953 to RET, and DE028397 to RET). The authors would also like to acknowledge Leah Melnikov and Adelle Melnikov for their assistance with image selection for analysis and Dr. Andrzej Fertala and Dr. Jolanta Fertala for their assistance in the acquisition and analysis of picrosirius red images.

## AUTHOR CONTRIBUTIONS

Conceptualization, R.E.T., A.C.; Investigation, A.C., E.M., A.C., and T.L.; Visualization, A.C.; Writing, A.C. and R.E.T.; Supervision, R.E.T.; Funding Acquisition, R.E.T.

## DATA AVAILABILITY STATEMENT

The data supporting this study is provided in the manuscript and supplement, with additional data available upon request.

## Notes

### Competing Interest Statement

The authors have declared no competing interest.

