## Supplemental Figures for "COX2-independent and COX2-dependent effects of naproxen on bone quality, osteocytes, and fatigue fracture healing in male and female mice"

### SUPPLEMENTAL DATA

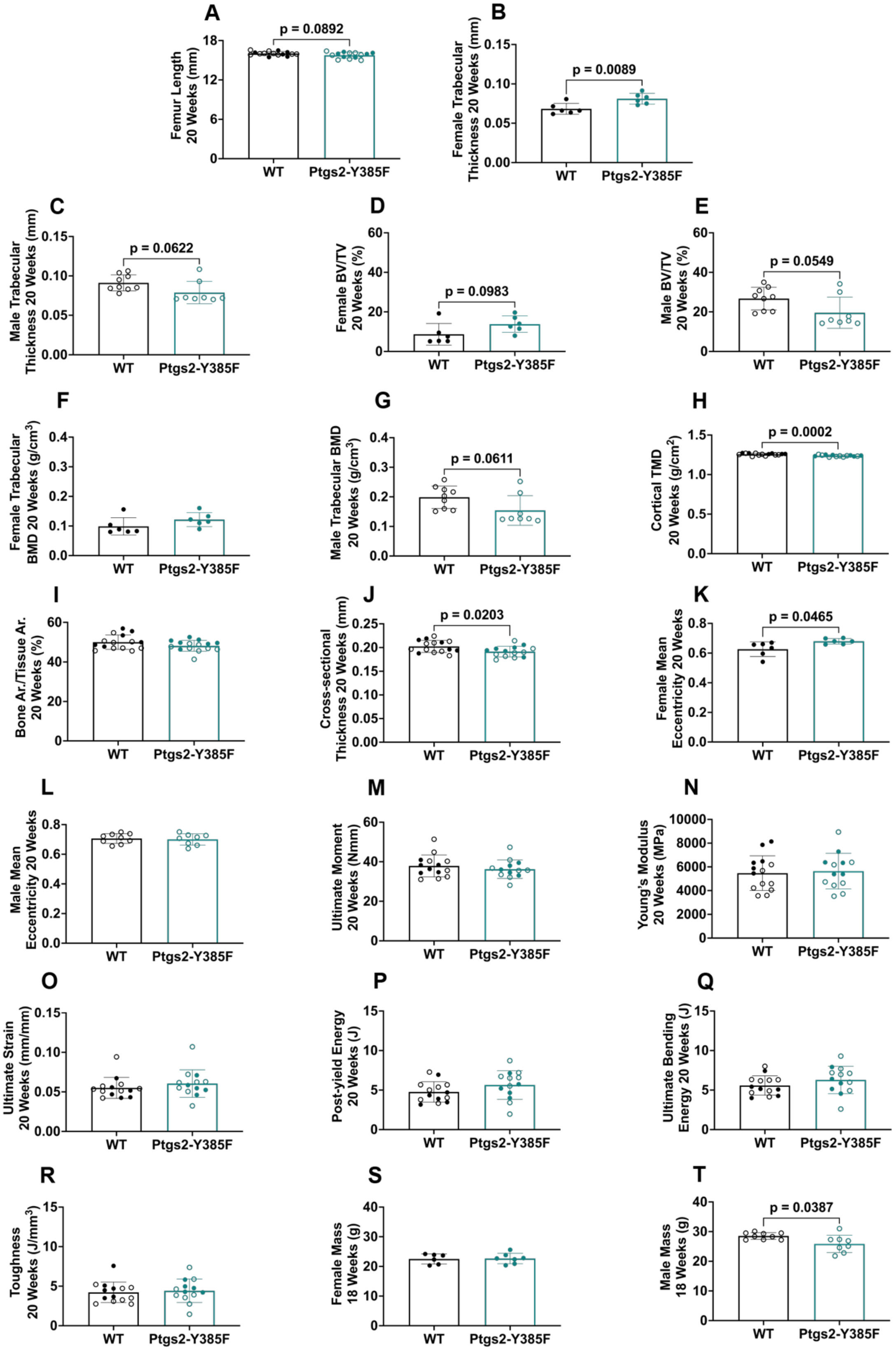

**Supplementary Figure 1: Ptgs2-Y385F mice are comparable to WT littermates in bone mechanical performance at 20 weeks of age, but display sexual dimorphism in measures of bone geometry.** Bone geometry was quantified in mice aged 20 weeks at experimental end using microCT scanning and reconstruction of non-loaded femurs and femoral mechanical performance was measured using standard three-point bending. Bone geometry parameters include (A) femur length, (B) female trabecular thickness, (C) male trabecular thickness, (D) female trabecular bone volume per tissue volume (BV/TV), (E) male BV/TV, (F) female trabecular bone mineral density (BMD), (G) male trabecular BMD, (H) cortical tissue mineral density (TMD), (I) cortical bone area per tissue area (Bone Ar./ Tissue Ar.), (J) cortical cross-sectional thickness, (K) female cortical mean eccentricity, and (L) male cortical mean eccentricity. n=6-7 females and n=8-9 males per group. Mechanical performance parameters include (M) ultimate moment, (N) Young's modulus, (O) ultimate strain, (P) post-yield energy, (Q) ultimate bending energy, and (R) toughness. n=13-14 per group. Body mass at 18 weeks is included as a measure of size at skeletal maturity in (S) female and (T) male mice. n=6-7 females and 8-9 males per group. A p value below 0.05 was considered significant and a p value below 0.1 was considered trending. Female replicates are displayed as filled circles and male replicates as empty circles.

| | **Number** | **Total Number** | **Percentage of Total Mice** | ***Expected Percentage*** |
|---|---|---|---|---|
| **Wild-Type:** | 108 | 335 | 32.2 | *25* |
| **Heterozygous:** | 182 | | 54.3 | *50* |
| **Ptgs2-Y385F:** | 45 | | 13.4 | *25* |
| **Males:** | 171 | 335 | 51.0 | *50* |
| **Females:** | 164 | | 49.0 | *50* |

**Supplementary Table 1: The Ptgs2-Y385F mutation does not follow expected Mendelian ratios for heterozygous x heterozygous breeding pairs.** Genotyping results from 335 total mice generated using heterozygous dams and bucks showing a decreased percentage of Ptgs2-Y385F mice as compared to the normal mendelian ratios published previously. The same group of mice was analyzed for the ratio of male and female mice and saw no difference from expected ratios.

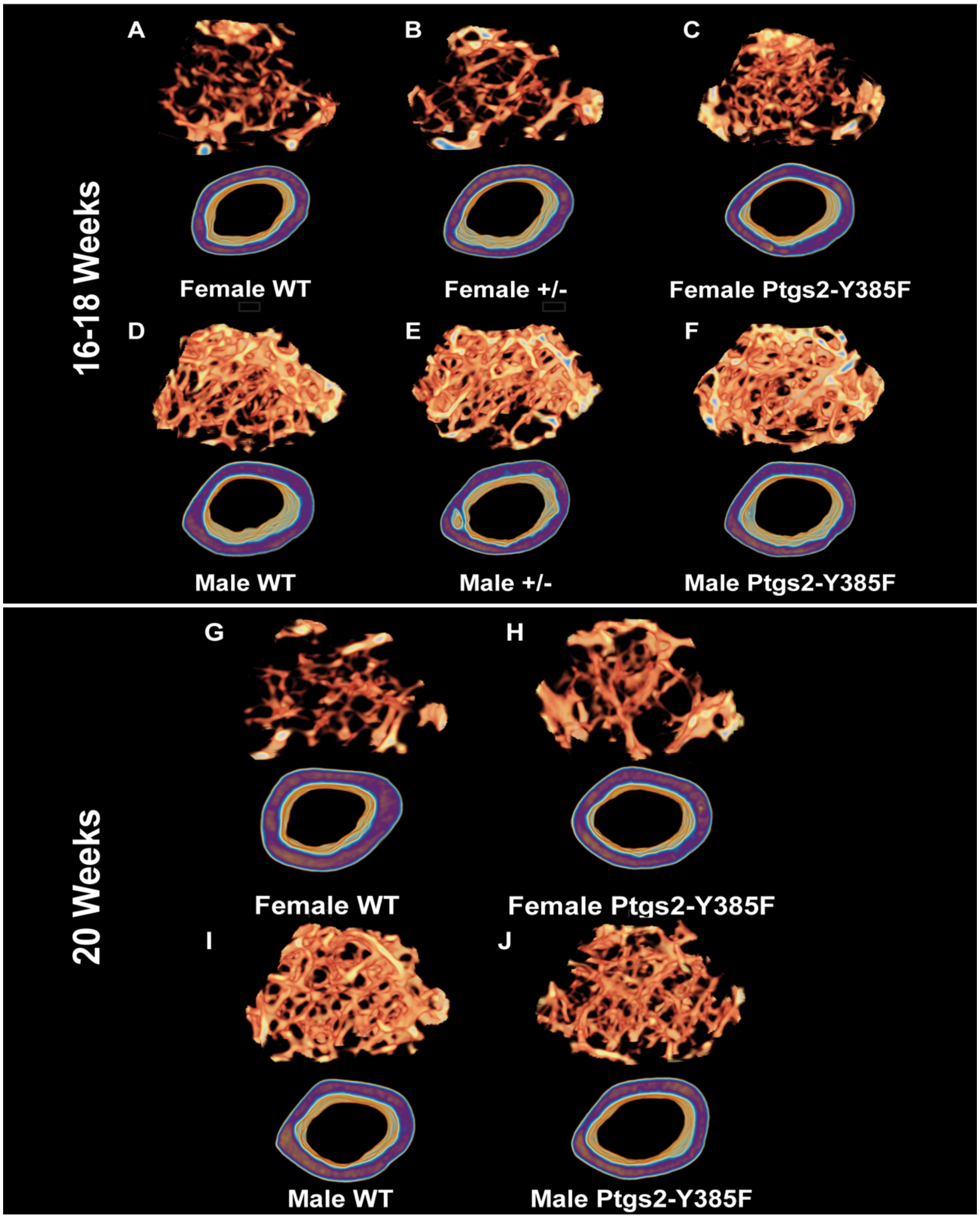


**Supplementary Figure 2: Representative microCT reconstructions of trabecular and cortical bone comparisons between WT and Ptgs2-Y385F mice at 16-18 weeks and 20 weeks old.** Quantifications of bone geometry are found in Figure 1 and Supplementary Figure 1.

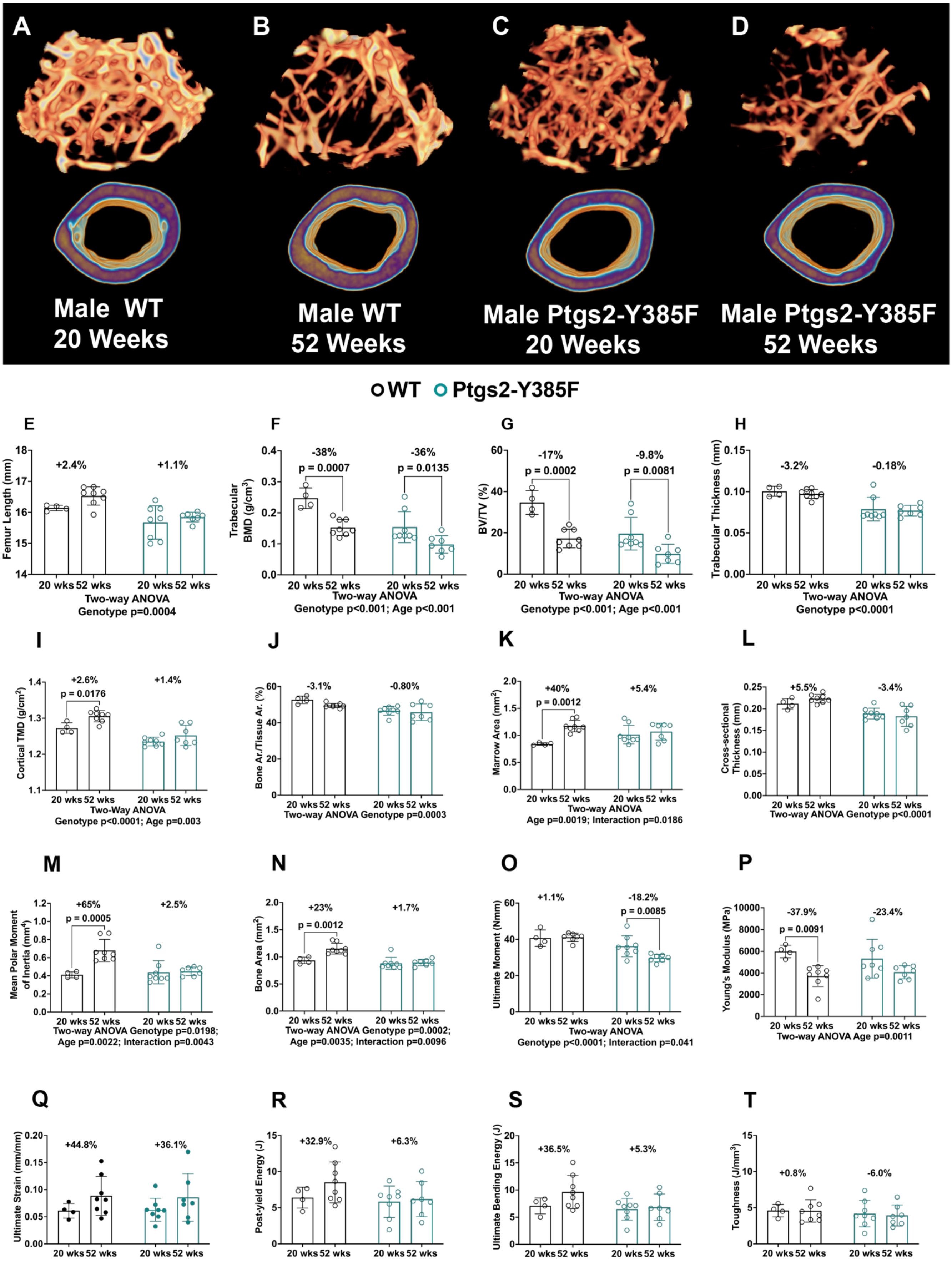

A
B
C
D
Male WT
20 Weeks
Male WT
52 Weeks
Male Ptgs2-Y385F
20 Weeks
Male Ptgs2-Y385F
52 Weeks
WT Ptgs2-Y385F
E
Femur Length (mm)
+2.4%
+1.1%
20 wks 52 wks 20 wks 52 wks
Two-way ANOVA
Genotype p=0.0004
F
Trabecular BMD (g/cm³)
-38%
p = 0.0007
-36%
p = 0.0135
Two-way ANOVA
Genotype p<0.001; Age p<0.001
G
BV/TV (%)
-17%
p = 0.0002
-9.8%
p = 0.0081
Two-way ANOVA
Genotype p<0.001; Age p<0.001
H
Trabecular Thickness (mm)
-3.2%
-0.18%
Two-way ANOVA
Genotype p<0.0001
I
Cortical TMD (g/cm³)
+2.6%
p = 0.0176
+1.4%
Two-Way ANOVA
Genotype p<0.0001; Age p=0.003
J
Bone Ar./Tissue Ar. (%)
-3.1%
-0.80%
Two-way ANOVA Genotype p=0.0003
K
Marrow Area (mm²)
+40%
p = 0.0012
+5.4%
Two-way ANOVA
Age p=0.0019; Interaction p=0.0186
L
Cross-sectional Thickness (mm)
+5.5%
-3.4%
Two-way ANOVA Genotype p<0.0001
M
Mean Polar Moment of Inertia (mm⁴)
+65%
p = 0.0005
+2.5%
Two-way ANOVA Genotype p=0.0198;
Age p=0.0022; Interaction p=0.0043
N
Bone Area (mm²)
+23%
p = 0.0012
+1.7%
Two-way ANOVA Genotype p=0.0002;
Age p=0.0035; Interaction p=0.0096
O
Ultimate Moment (Nmm)
+1.1%
-18.2%
p = 0.0085
Two-way ANOVA
Genotype p<0.0001; Interaction p=0.041
P
Young's Modulus (MPa)
-37.9%
p = 0.0091
-23.4%
Two-way ANOVA Age p=0.0011
Q
Ultimate Strain (mm/mm)
+44.8%
+36.1%
R
Post-yield Energy (J)
+32.9%
+6.3%
S
Ultimate Bending Energy (J)
+36.5%
+5.3%
T
Toughness (J/mm³)
+0.8%
-6.0%

**Supplementary Figure 3: Ptgs2-Y385F males exhibit expected changes to trabecular bone geometry with aging, but altered cortical bone geometry changes leading to decreased mechanical performance in comparison to WT, shared genetic background, males.** Bone geometry was quantified using microCT scanning and reconstruction of non-loaded femurs from WT males and Ptgs2-Y385F males aged 20 and 52 weeks, and femoral mechanical performance was measured using standard three-point bending. (A-D) Representative microCT reconstructions of trabecular and cortical bone regions for male WT and male Ptgs2-Y385F mice. Bone geometry parameters include (E) femur length, (F) trabecular bone mineral density (BMD), (G) trabecular bone volume per tissue volume (BV/TV), (H) trabecular thickness, (I) cortical tissue mineral density (TMD), (J) cortical bone area per tissue area (Bone Ar./ Tissue Ar.), (K) marrow area, (L) cortical cross-sectional thickness, (M) mean polar moment of inertia, and (N) bone area. Mechanical performance parameters include (O) ultimate moment, (P) Young's modulus, (Q) ultimate strain, (R) post-yield energy, (S) ultimate bending energy, and (T) toughness. n=4-8 per group. A p value below 0.05 was considered significant and a p value below 0.1 was considered trending. Female replicates are displayed as filled circles and male replicates as empty circles.

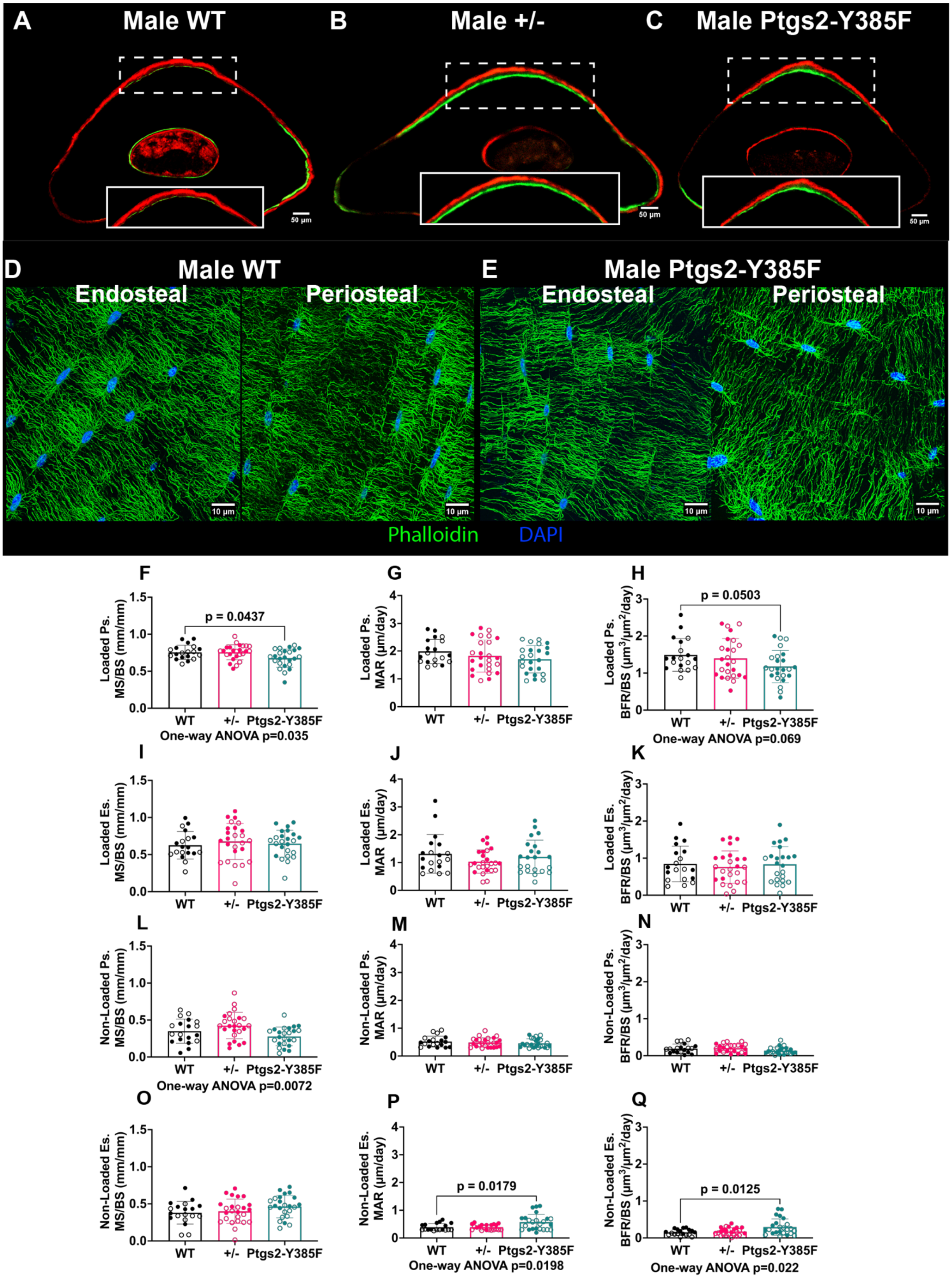

A Male WT
B Male +/-
C Male Ptgs2-Y385F
50 μm
D Male WT
E Male Ptgs2-Y385F
Endosteal
Periosteal
10 μm
Phalloidin DAPI
F Loaded Ps. MS/BS (mm/mm)
p = 0.0437
One-way ANOVA p=0.035
G Loaded Ps. MAR (μm/day)
H Loaded Ps. BFR/BS (μm3/μm2/day)
p = 0.0503
One-way ANOVA p=0.069
I Loaded Es. MS/BS (mm/mm)
J Loaded Es. MAR (μm/day)
K Loaded Es. BFR/BS (μm3/μm2/day)
L Non-Loaded Ps. MS/BS (mm/mm)
One-way ANOVA p=0.0072
M Non-Loaded Ps. MAR (μm/day)
N Non-Loaded Ps. BFR/BS (μm3/μm2/day)
O Non-Loaded Es. MS/BS (mm/mm)
P Non-Loaded Es. MAR (μm/day)
p = 0.0179
One-way ANOVA p=0.0198
Q Non-Loaded Es. BFR/BS (μm3/μm2/day)
p = 0.0125
One-way ANOVA p=0.022
WT +/- Ptgs2-Y385F

**Supplementary Figure 4: Changes in load-induced bone formation caused by the Ptgs2-Y385F mutation are predominantly seen in the periosteal mineralizing surfaces of loaded limbs and basal state (non-loaded) endosteal surface remodeling.** Bone formation in response to uniaxial forelimb compression was measured in ulnar mid-diaphysis sections using dynamic histomorphometry with calcein (green) and alizarin red (red) bone formation labels in mice aged 16-18 weeks at experiment end. Osteocyte dendritic networks were visualized in mid-cortical sections of non-loaded femurs using phalloidin F-actin staining (green) and DAPI nuclear staining (blue), and networks were quantified using dendrite counting near the nucleus (peri-nuclear) and at a standard distance away from the nucleus where dendrites commonly ended (terminal) and dendritic density was measured using the moving band method in mice aged 20 weeks at experiment end. (A-C) Representative sections from male WT, heterozygous (+/-), and Ptgs2-Y385F mice. (D-E) Representative sections from male WT and Ptgs2-Y385F mice showing the dendritic network near the endosteal and the periosteal surfaces of the bone. Measured dynamic histomorphometry outcomes included (F) loaded periosteal mineralizing surface per bone surface (Ps. MS/BS), (G) loaded periosteal mineral apposition rate (Ps. MAR), (H) loaded periosteal bone formation rate per bone surface (Ps. BFR/BS), (I-K) corresponding measurements on the endosteal surface, (L) non-loaded periosteal mineralizing surface per bone surface (Ps. MS/BS), (M) non-loaded periosteal mineral apposition rate (Ps. MAR), (N) non-loaded periosteal bone formation rate per bone surface (Ps. BFR/BS), and (O-Q) corresponding measurements on the endosteal surface. n=19-25 per group. Quantifications of dendrite number and density are found in Figure 2. A p value below 0.05 was considered significant and a p value below 0.1 was considered trending. Female replicates are displayed as filled circles and male replicates as empty circles.

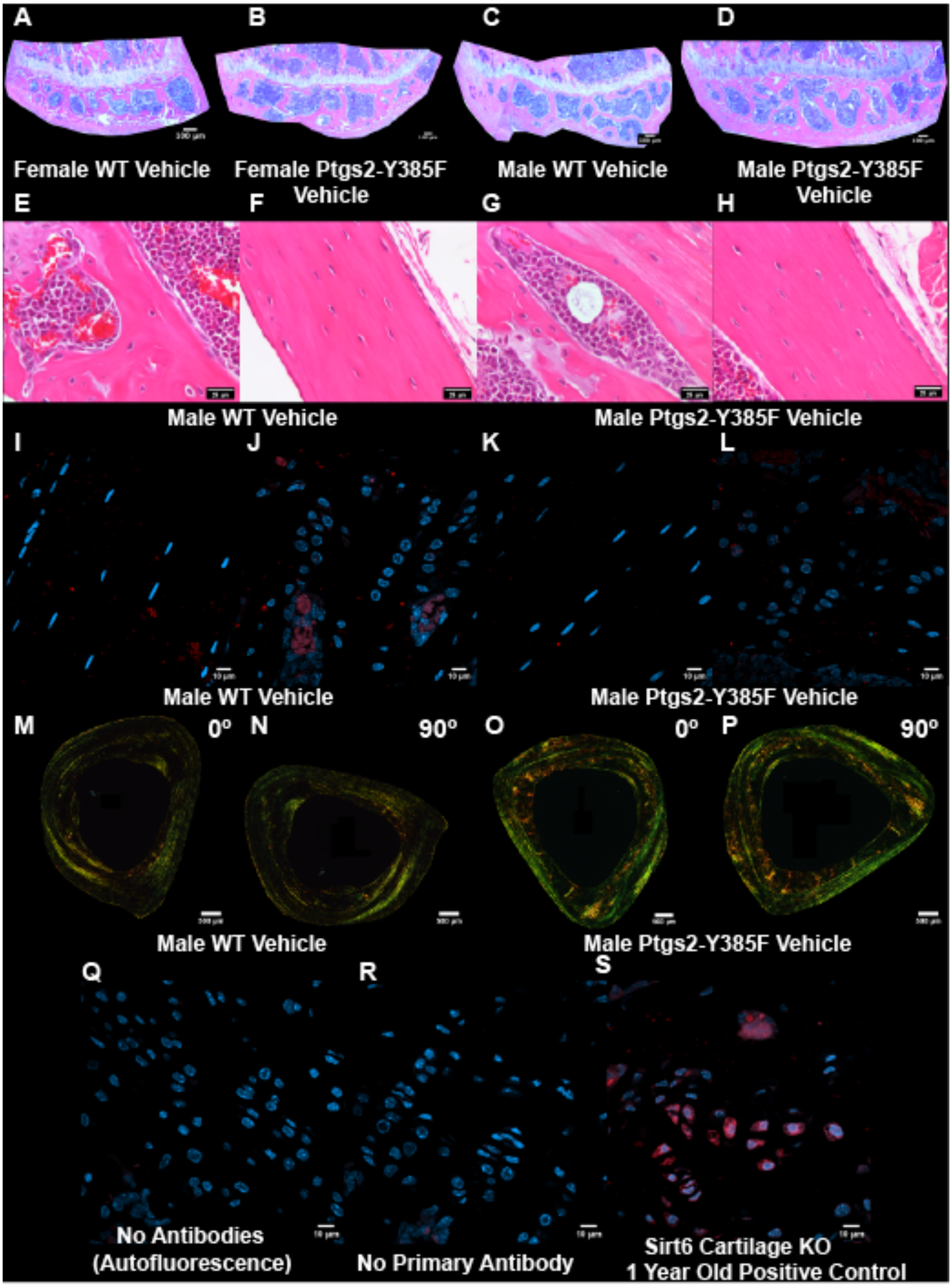
A
B
C
D
Female WT Vehicle
Female Ptgs2-Y385F Vehicle
Male WT Vehicle
Male Ptgs2-Y385F Vehicle
E
F
G
H
Male WT Vehicle
Male Ptgs2-Y385F Vehicle
I
J
K
L
Male WT Vehicle
Male Ptgs2-Y385F Vehicle
M
0º
N
90º
O
0º
P
90º
Male WT Vehicle
Male Ptgs2-Y385F Vehicle
Q
R
S
No Antibodies (Autofluorescence)
No Primary Antibody
Sirt6 Cartilage KO 1 Year Old Positive Control

**Supplementary Figure 5: Representative histological images of tibial growth plates across both sexes and genotypes, male osteoblast and osteocyte lacunae images across genotypes, male MMP-13 expression in bone and growth plate chondrocytes across genotypes, male picrosirius red images across genotypes, and immunofluorescence controls.** (A-P) Representative images of histology sections for females and males not included in Figure 3. Corresponding quantifications of histology are found in Figure 3. Immunofluorescence controls included (Q) a no primary or secondary antibody control to visualize tissue autofluorescence, (R) a no primary antibody control to ensure the specificity of the secondary antibody, and (S) a positive control of a joint from a Sirt6 cartilage-specific knockout mouse aged one year exhibiting extensive MMP-13 expression in growth plate chondrocytes due to accelerated age-induced osteoarthritis and cartilage damage.

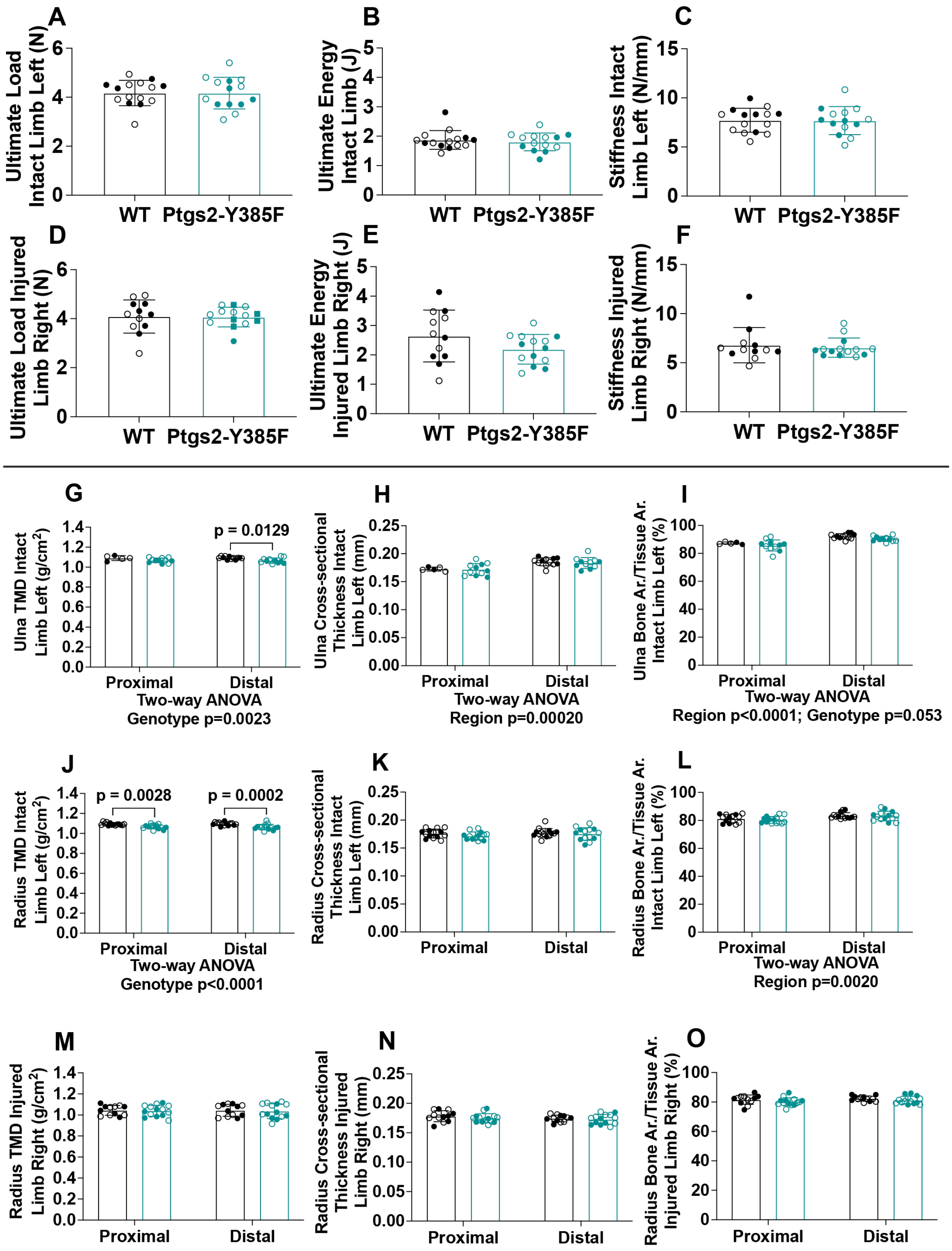
A
Ultimate Load Intact Limb Left (N)
WT
Ptgs2-Y385F
B
Ultimate Energy Intact Limb (J)
WT
Ptgs2-Y385F
C
Stiffness Intact Limb Left (N/mm)
WT
Ptgs2-Y385F
D
Ultimate Load Injured Limb Right (N)
WT
Ptgs2-Y385F
E
Ultimate Energy Injured Limb Right (J)
WT
Ptgs2-Y385F
F
Stiffness Injured Limb Right (N/mm)
WT
Ptgs2-Y385F
G
Ulna TMD Intact Limb Left (g/cm2)
p = 0.0129
Proximal
Distal
Two-way ANOVA
Genotype p=0.0023
H
Ulna Cross-sectional Thickness Intact Limb Left (mm)
Proximal
Distal
Two-way ANOVA
Region p=0.00020
I
Ulna Bone Ar./Tissue Ar. Intact Limb Left (%)
Proximal
Distal
Two-way ANOVA
Region p<0.0001; Genotype p=0.053
J
Radius TMD Intact Limb Left (g/cm2)
p = 0.0028
p = 0.0002
Proximal
Distal
Two-way ANOVA
Genotype p<0.0001
K
Radius Cross-sectional Thickness Intact Limb Left (mm)
Proximal
Distal
L
Radius Bone Ar./Tissue Ar. Intact Limb Left (%)
Proximal
Distal
Two-way ANOVA
Region p=0.0020
M
Radius TMD Injured Limb Right (g/cm2)
Proximal
Distal
N
Radius Cross-sectional Thickness Injured Limb Right (mm)
Proximal
Distal
O
Radius Bone Ar./Tissue Ar. Injured Limb Right (%)
Proximal
Distal

**Supplementary Figure 6: Fatigue fracture limbs 15 days post-injury of WT and Ptgs2-Y385F mice show comparable mechanical performance under compression and minimal difference in radial or ulnar bone geometry across genotypes.** Mechanical performance of the bone callus and the intact limb were measured separately using monotonic compression to failure. Bone geometry was quantified using microCT scanning and reconstruction of intact and injured forelimbs to quantify genotype-driven differences in radial and ulnar bone geometry. Quantifications in the intact (left) limb of (A) ultimate load, (B) ultimate energy, and (C) stiffness. (D-F) Corresponding measures in injured limbs. N=13-14 per group. Quantifications in the intact (left) limb of (G) ulnar tissue mineral density (TMD), (H) ulnar cortical cross-sectional thickness, (I) ulnar bone area per tissue area (Bone Ar./Tissue Ar.). n=5-12 per group. Quantifications in the intact (left) limb of radial TMD, (J) radial cortical cross-sectional thickness, (K) radial Bone Ar./Tissue Ar. (M-O) Corresponding radial bone geometry measures from the injured (right) limb. n=12-14 per group. A p value below 0.05 was considered significant and a p value below 0.1 was considered trending. Female replicates are displayed as filled circles and male replicates as empty circles.

| Treatment | High Dose Naproxen Sodium (41.6 mg/kg/day) | Low Dose Naproxen Sodium (10.9 mg/kg/day) |
|---|---|---|
| Average mouse mass (g) Week 1 | 21.9 | - |
| Average mouse mass (g) Week 2 | 22.9 | - |
| Average mouse mass (g) Week 3 | 22.4 | - |
| Average mouse mass (g) Week 4 | 23.1 | 23.1 |
| Average Water Consumed/mouse/day (mL) Check 1 | 4.3 | 6.0 |
| Average Water Consumed/mouse/day (mL) Check 2 | 4.1 | - |
| Average Water Consumed/mouse/day (mL) Check 3 | 4.3 | - |
| Average Water Consumed/mouse/day (mL) Check 4 | 4.7 | - |

**Supplementary Table 2: High dose naproxen drinking water has comparable consumption patterns to low dose naproxen drinking water by C57BL6/J females and does not significantly change body mass over 30 days of treatment.** Average mouse body mass and drinking water consumption per day was calculated for mice receiving high dose naproxen sodium (41.6 mg/kg/day) and compared to a measurement of mice receiving low dose naproxen sodium (10.9 mg/kg/day). This was done to ensure that changes in water taste from increased dosage did not cause deviations from the calculated body mass and water consumption used to calculate dosing.

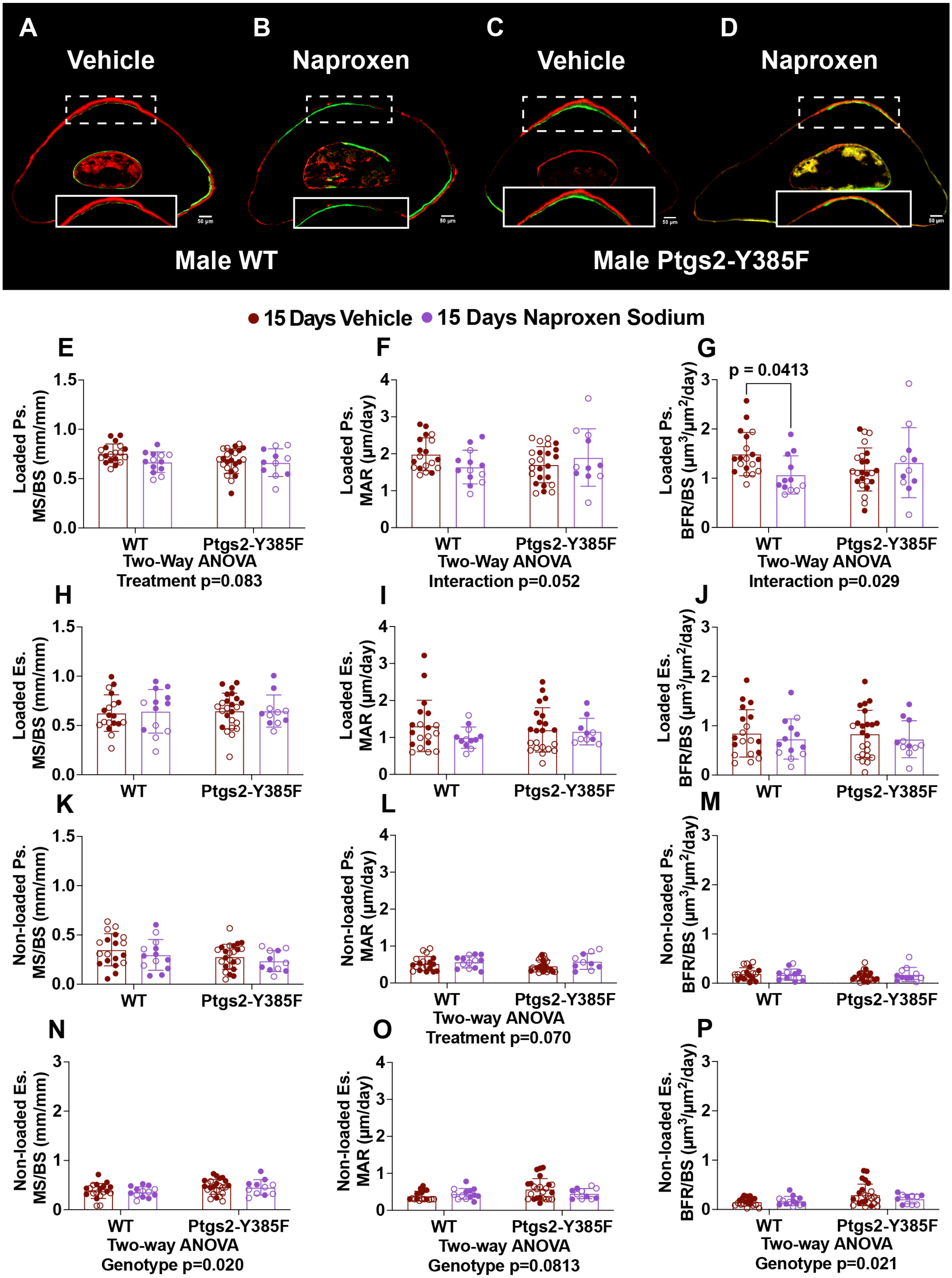

A
Vehicle
B
Naproxen
C
Vehicle
D
Naproxen
Male WT
Male Ptgs2-Y385F
15 Days Vehicle
15 Days Naproxen Sodium
E
Loaded Ps. MS/BS (mm/mm)
WT
Ptgs2-Y385F
Two-Way ANOVA
Treatment p=0.083
F
Loaded Ps. MAR (μm/day)
WT
Ptgs2-Y385F
Two-Way ANOVA
Interaction p=0.052
G
p = 0.0413
Loaded Ps. BFR/BS (μm3/μm2/day)
WT
Ptgs2-Y385F
Two-Way ANOVA
Interaction p=0.029
H
Loaded Es. MS/BS (mm/mm)
WT
Ptgs2-Y385F
I
Loaded Es. MAR (μm/day)
WT
Ptgs2-Y385F
J
Loaded Es. BFR/BS (μm3/μm2/day)
WT
Ptgs2-Y385F
K
Non-loaded Ps. MS/BS (mm/mm)
WT
Ptgs2-Y385F
L
Non-loaded Ps. MAR (μm/day)
WT
Ptgs2-Y385F
Two-way ANOVA
Treatment p=0.070
M
Non-loaded Ps. BFR/BS (μm3/μm2/day)
WT
Ptgs2-Y385F
N
Non-loaded Es. MS/BS (mm/mm)
WT
Ptgs2-Y385F
Two-way ANOVA
Genotype p=0.020
O
Non-loaded Es. MAR (μm/day)
WT
Ptgs2-Y385F
Two-way ANOVA
Genotype p=0.0813
P
Non-loaded Es. BFR/BS (μm3/μm2/day)
WT
Ptgs2-Y385F
Two-way ANOVA
Genotype p=0.021

**Supplementary Figure 7: Naproxen treatment for 15 days decreases load-induced bone formation predominantly through decreased mineral apposition rate on the periosteal surface of the loaded limb and does not affect endosteal bone remodeling.** Bone formation in response to uniaxial forelimb compression was measured in ulnar mid-diaphysis sections using dynamic histomorphometry with calcein (green) and alizarin red (red) bone formation labels in mice aged 16-18 weeks at experiment end who received vehicle or naproxen drinking water for 15 days. (A-D) Representative sections from male WT and Ptgs2-Y385F mice treated for 15 days. Measured dynamic histomorphometry outcomes included (E) loaded periosteal mineralizing surface per bone surface (Ps. MS/BS), (F) loaded periosteal mineral apposition rate (Ps. MAR), (G) loaded periosteal bone formation rate per bone surface (Ps. BFR/BS), (H-J) corresponding measurements on the endosteal surface, (K) non-loaded periosteal mineralizing surface per bone surface (Ps. MS/BS), (L) non-loaded periosteal mineral apposition rate (Ps. MAR), (M) non-loaded periosteal bone formation rate per bone surface (Ps. BFR/BS), and (N-P) corresponding measurements on the endosteal surface. n=19-25 per group. Quantifications of dendrite number and density are found in Figure 2. A p value below 0.05 was considered significant and a p value below 0.1 was considered trending. Female replicates are displayed as filled circles and male replicates as empty circles.

| Treatment Group | # Samples with Ps. Double Labels | # Samples with Single Ps. Labels | # Samples without Ps. Labels | # Samples assigned 0.3 Ps. MAR | # Samples with Es. Double Labels | # Samples with Single Es. Labels | # Samples without Es. Labels | # Samples assigned 0.3 Es. MAR |
|---|---|---|---|---|---|---|---|---|
| Vehicle WT | 71 | 68 | 0 | 5 | 52 | 69 | 0 | 24 |
| Vehicle Heterozygous | 95 | 100 | 0 | 6 | 79 | 95 | 0 | 21 |
| Vehicle Ptgs2-Y385F | 90 | 102 | 0 | 14 | 81 | 95 | 0 | 19 |
| Naproxen Sodium WT | 50 | 59 | 0 | 9 | 40 | 54 | 0 | 12 |
| Naproxen Sodium Ptgs2-Y385F | 41 | 43 | 0 | 6 | 41 | 50 | 0 | 10 |

**Supplementary Table 3: Tabular description of dynamic histomorphometry results.** The number of technical replicates within each treatment group that had double labeling, single labeling, no labeling, and assumed an assumed mineral apposition rate (MAR) of 0.03 to account for samples without double labeling on both the periosteal and endosteal surface are listed.

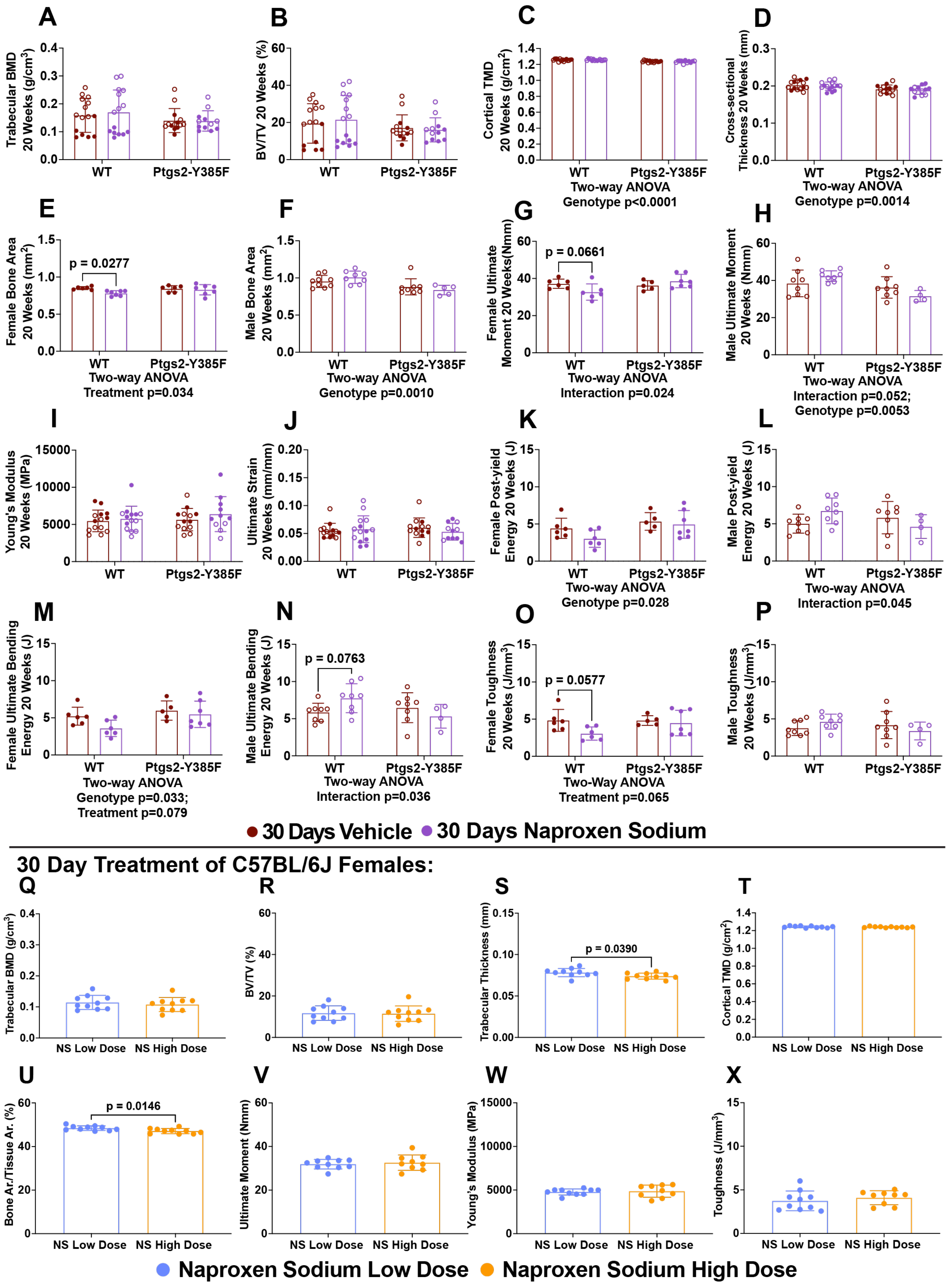
A
Trabecular BMD 20 Weeks (g/cm³)
WT
Ptgs2-Y385F
B
BV/TV 20 Weeks (%)
WT
Ptgs2-Y385F
C
Cortical TMD 20 Weeks (g/cm²)
WT
Ptgs2-Y385F
Two-way ANOVA
Genotype p<0.0001
D
Cross-sectional Thickness 20 Weeks (mm)
WT
Ptgs2-Y385F
Two-way ANOVA
Genotype p=0.0014
E
Female Bone Area 20 Weeks (mm²)
p = 0.0277
WT
Ptgs2-Y385F
Two-way ANOVA
Treatment p=0.034
F
Male Bone Area 20 Weeks (mm²)
WT
Ptgs2-Y385F
Two-way ANOVA
Genotype p=0.0010
G
Female Ultimate Moment 20 Weeks(Nmm)
p = 0.0661
WT
Ptgs2-Y385F
Two-way ANOVA
Interaction p=0.024
H
Male Ultimate Moment 20 Weeks (Nmm)
WT
Ptgs2-Y385F
Two-way ANOVA
Interaction p=0.052;
Genotype p=0.0053
I
Young's Modulus 20 Weeks (MPa)
WT
Ptgs2-Y385F
J
Ultimate Strain 20 Weeks (mm/mm)
WT
Ptgs2-Y385F
K
Female Post-yield Energy 20 Weeks (J)
WT
Ptgs2-Y385F
Two-way ANOVA
Genotype p=0.028
L
Male Post-yield Energy 20 Weeks (J)
WT
Ptgs2-Y385F
Two-way ANOVA
Interaction p=0.045
M
Female Ultimate Bending Energy 20 Weeks (J)
WT
Ptgs2-Y385F
Two-way ANOVA
Genotype p=0.033;
Treatment p=0.079
N
Male Ultimate Bending Energy 20 Weeks (J)
p = 0.0763
WT
Ptgs2-Y385F
Two-way ANOVA
Interaction p=0.036
O
Female Toughness 20 Weeks (J/mm³)
p = 0.0577
WT
Ptgs2-Y385F
Two-Way ANOVA
Treatment p=0.065
P
Male Toughness 20 Weeks (J/mm³)
WT
Ptgs2-Y385F
30 Days Vehicle
30 Days Naproxen Sodium
30 Day Treatment of C57BL/6J Females:
Q
Trabecular BMD (g/cm³)
NS Low Dose
NS High Dose
R
BV/TV (%)
NS Low Dose
NS High Dose
S
Trabecular Thickness (mm)
p = 0.0390
NS Low Dose
NS High Dose
T
Cortical TMD (g/cm²)
NS Low Dose
NS High Dose
U
Bone Ar./Tissue Ar. (%)
p = 0.0146
NS Low Dose
NS High Dose
V
Ultimate Moment (Nmm)
NS Low Dose
NS High Dose
W
Young's Modulus (MPa)
NS Low Dose
NS High Dose
X
Toughness (J/mm³)
NS Low Dose
NS High Dose
Naproxen Sodium Low Dose
Naproxen Sodium High Dose

**Supplementary Figure 8: Naproxen's effects on bone mechanical performance are likely related more to extended use than dosage when used within the daily recommended dose.** Bone geometry was quantified using microCT scanning and reconstruction of non-loaded femurs from mice treated with vehicle or naproxen for 30 days and femoral mechanical performance was measured using standard three-point bending. In a separate cohort, female C57BL/6J received low dose (10.9 mg/kg/day) naproxen (the same dose as all other naproxen treatment group in this study), or high dose (41.6 mg/kg/day) naproxen for 30 days. Bone geometry and mechanical performance of their non-loaded femurs were measured as above. Bone geometry parameters include (A) trabecular bone mineral density (BMD), (B) trabecular bone volume per tissue volume (BV/TV), (C) cortical tissue mineral density (TMD), and (D) cortical cross-sectional thickness, and cortical bone area for (E) female and (F) male mice. n=14-15 females and n=12-15 males per group. Mechanical performance parameters include ultimate moment for (G) female and (H) male mice, (I) Young's modulus, (J) ultimate strain, post-yield energy for (K) female and (L) male mice, ultimate bending energy for (M) female and (N) male mice, and toughness for (O) female and (P) male mice. n=5-7 females and n=4-8 males per group. For C57BL/6J females, Bone geometry parameters include (Q) trabecular BMD, (R) trabecular BV/TV, (S) trabecular thickness, (T) cortical TMD, and (U) cortical Bone Ar./Tissue Ar. Mechanical performance parameters include (V) ultimate moment, (W) Young's modulus, and (X) toughness. n=9-10 per group. A p value below 0.05 was considered significant and a p value below 0.1 was considered trending. Female replicates are displayed as filled circles and male replicates as empty circles.

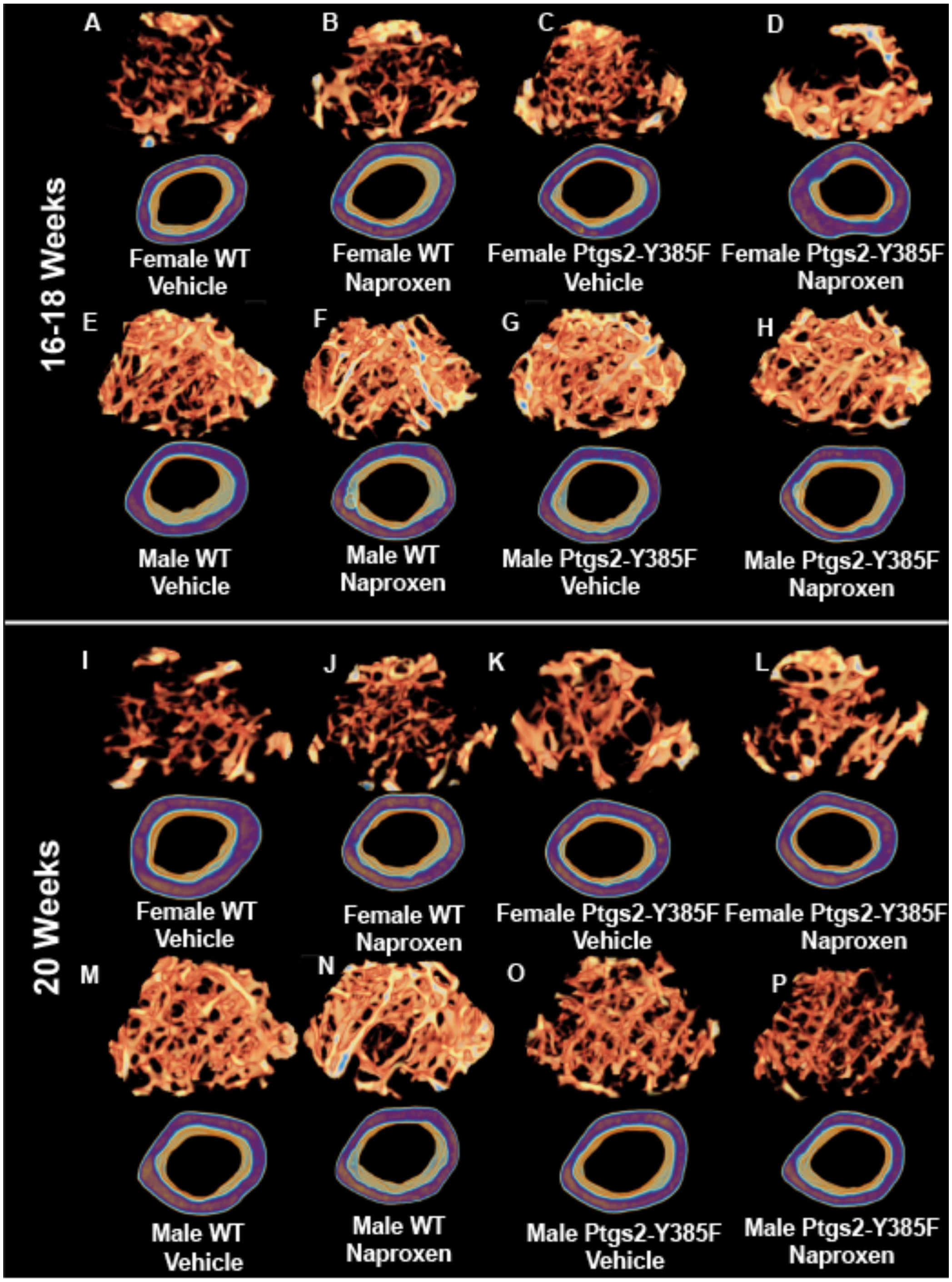


**Supplementary Figure 9: Representative microCT reconstructions of trabecular and cortical bone comparisons between WT and Ptgs2-Y385F mice after 15 days and 30 days of naproxen treatment.** Quantifications of bone geometry are found in Figure 5 and Supplementary Figure 7.

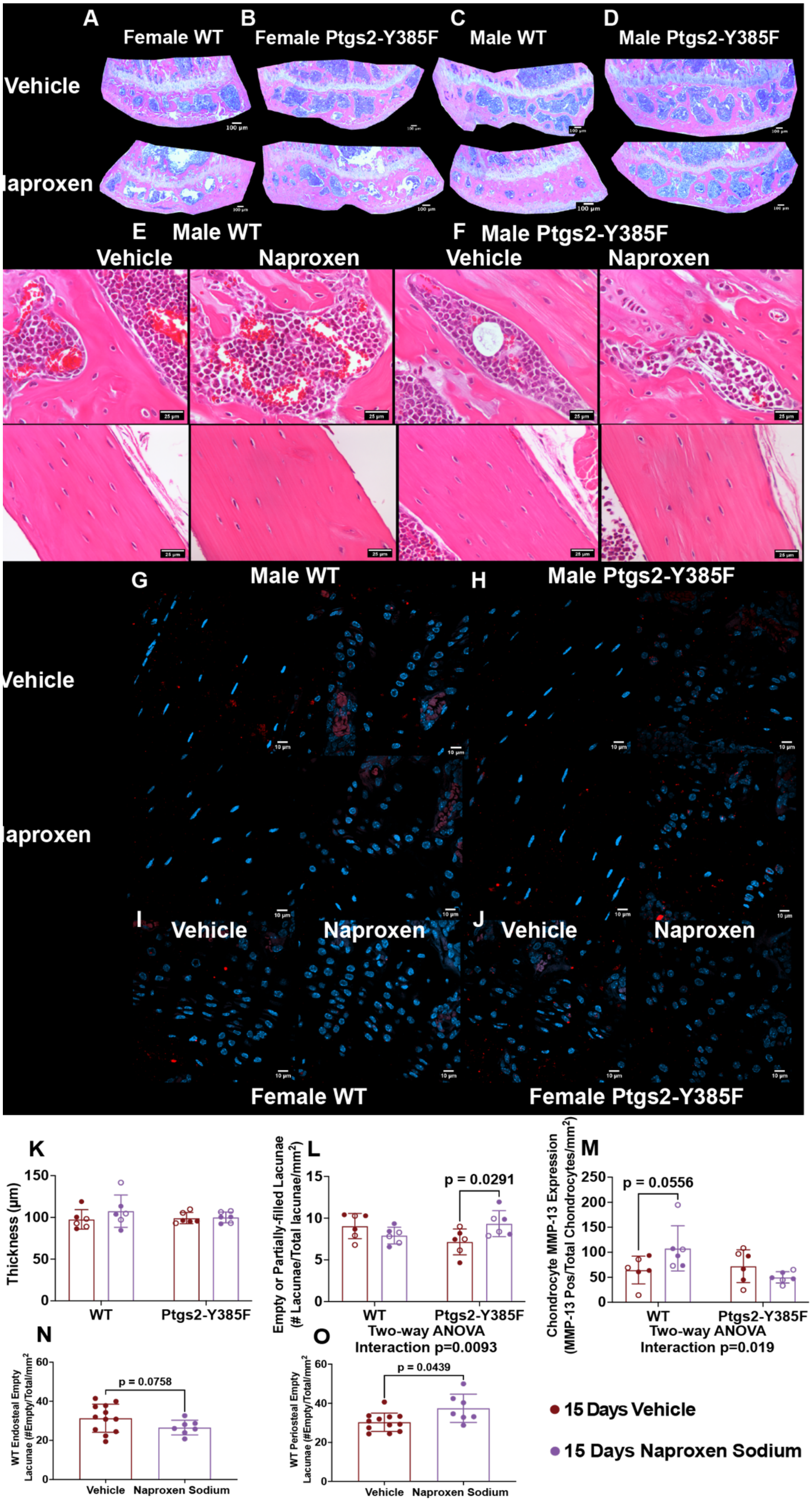

A
Female WT
B
Female Ptgs2-Y385F
C
Male WT
D
Male Ptgs2-Y385F
Vehicle
Naproxen
E
Male WT
Vehicle
Naproxen
F
Male Ptgs2-Y385F
Vehicle
Naproxen
G
Male WT
H
Male Ptgs2-Y385F
Vehicle
Naproxen
I
Vehicle
Naproxen
J
Vehicle
Naproxen
Female WT
Female Ptgs2-Y385F
K
Thickness (µm)
WT
Ptgs2-Y385F
L
Empty or Partially-filled Lacunae (# Lacunae/Total lacunae/mm2)
p = 0.0291
WT
Ptgs2-Y385F
Two-way ANOVA
Interaction p=0.0093
M
Chondrocyte MMP-13 Expression (MMP-13 Pos/Total Chondrocytes/mm2)
p = 0.0556
WT
Ptgs2-Y385F
Two-way ANOVA
Interaction p=0.019
N
WT Endosteal Empty Lacunae (#Empty/Total/mm2
p = 0.0758
Vehicle
Naproxen Sodium
O
WT Periosteal Empty Lacunae (#Empty/Total/mm2
p = 0.0439
Vehicle
Naproxen Sodium
15 Days Vehicle
15 Days Naproxen Sodium

**Supplementary Figure 10: Naproxen treatment for 15 days does not affect tibial growth plate thickness in either sex or genotype, but alters osteocyte viability and growth plate chondrocyte MMP-13 expression differently between WT and Ptgs2-Y385F mice.** (A-J) Representative images of histology sections for females and males treated with vehicle or naproxen for 15 days and not included in Figure 6. Corresponding quantifications of histology are found in Figure 6. Quantifications of (K) tibial growth plate thickness, (L) the number of empty and partially-filled osteocyte lacunae in cortical bone, (M) MMP-13 expression in growth plate chondrocytes, and empty osteocyte lacunae (no DAPI staining) in dynamic histomorphometry sections near the (N) endosteal and (O) periosteal surfaces of the bone. A p value below 0.05 was considered significant and a p value below 0.1 was considered trending. Female replicates are displayed as filled circles and male replicates as empty circles.

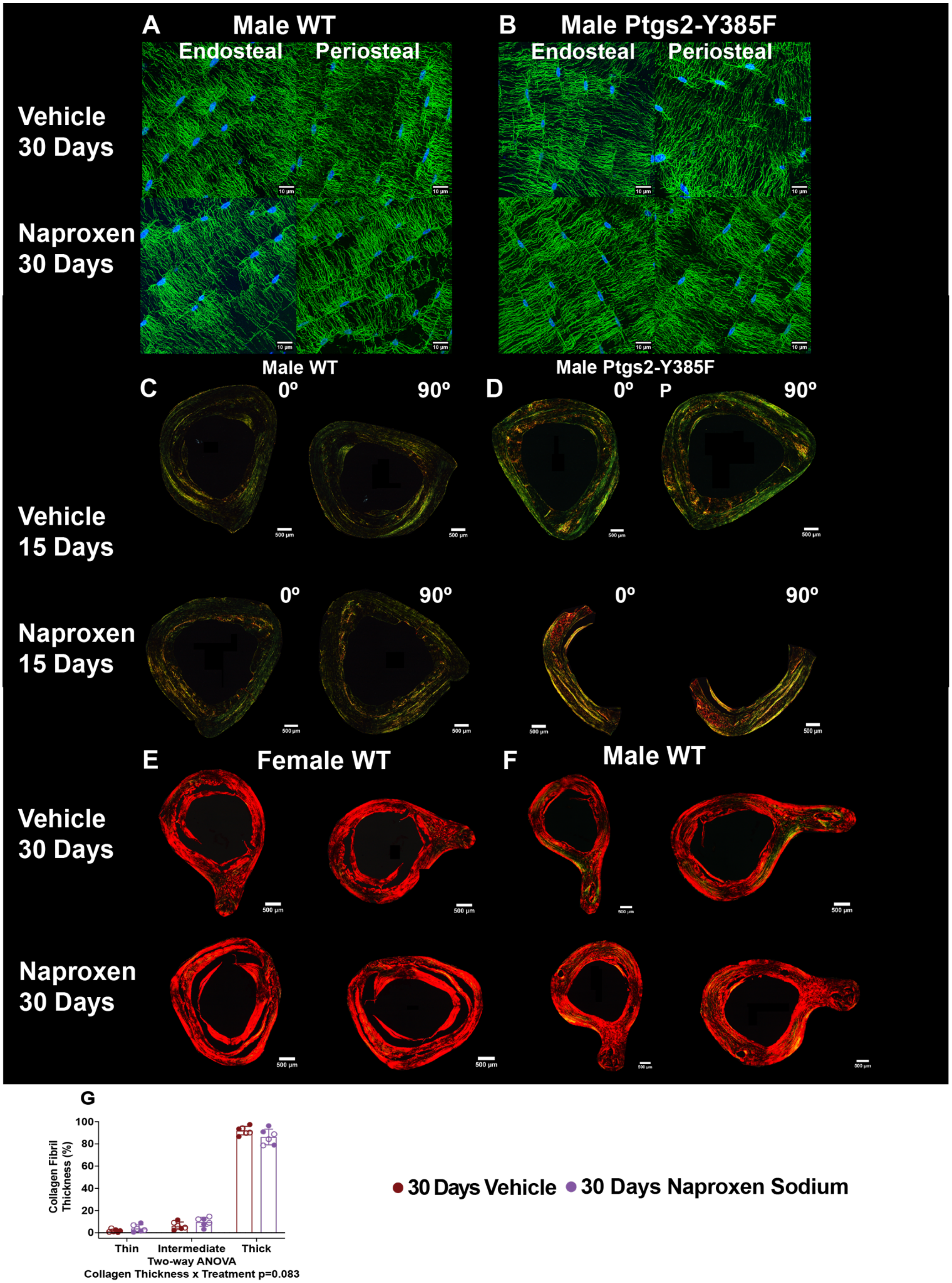
A
Male WT
Endosteal
Periosteal
B
Male Ptgs2-Y385F
Endosteal
Periosteal
Vehicle 30 Days
Naproxen 30 Days
Male WT
Male Ptgs2-Y385F
C
0º
90º
D
0º
P
90º
Vehicle 15 Days
Naproxen 15 Days
E
Female WT
F
Male WT
Vehicle 30 Days
Naproxen 30 Days
G
Collagen Fibril Thickness (%)
Thin
Intermediate
Thick
Two-way ANOVA
Collagen Thickness x Treatment p=0.083
30 Days Vehicle
30 Days Naproxen Sodium

**Supplementary Figure 11: Naproxen treatment for 30 days alters collagen fibril thickness similarly to 15 days of treatment in WT mice and does not correlate directly with bone toughness measurements.** Osteocyte dendritic networks were visualized in mid-cortical sections of non-loaded femurs using phalloidin F-actin staining (green) and DAPI nuclear staining (blue) in mice treated with vehicle or naproxen for 30 days, collagen fibril thickness was visualized using picrosirius red staining in mice treated for 15 or 30 days with vehicle or naproxen. (A-B) Representative sections from male WT and Ptgs2-Y385F mice treated for 30 days showing the dendritic network near the endosteal and the periosteal surfaces of the bone. (C-D) Representative images of tibial cross sections stained with picrosirius red from male WT and Ptgs2-Y385F mice aged 16-18 weeks at experiment end that received vehicle or naproxen for 15 days. (E-F) Representative images of tibial cross sections stained with picrosirius red from female and male WT mice aged 20 weeks at experiment end that received vehicle or naproxen for 30 days. (G) Quantification of thin (green), intermediate (yellow), and thick (red) collagen fibrils measured percentage of stained area in the cross section. n=6 per group. Differences in picrosirius red staining appearance are due to 15 day samples being embedded in paraffin and 30 day samples being embedded in OCT. A p value below 0.05 was considered significant and a p value below 0.1 was considered trending. Female replicates are displayed as filled circles and male replicates as empty circles.

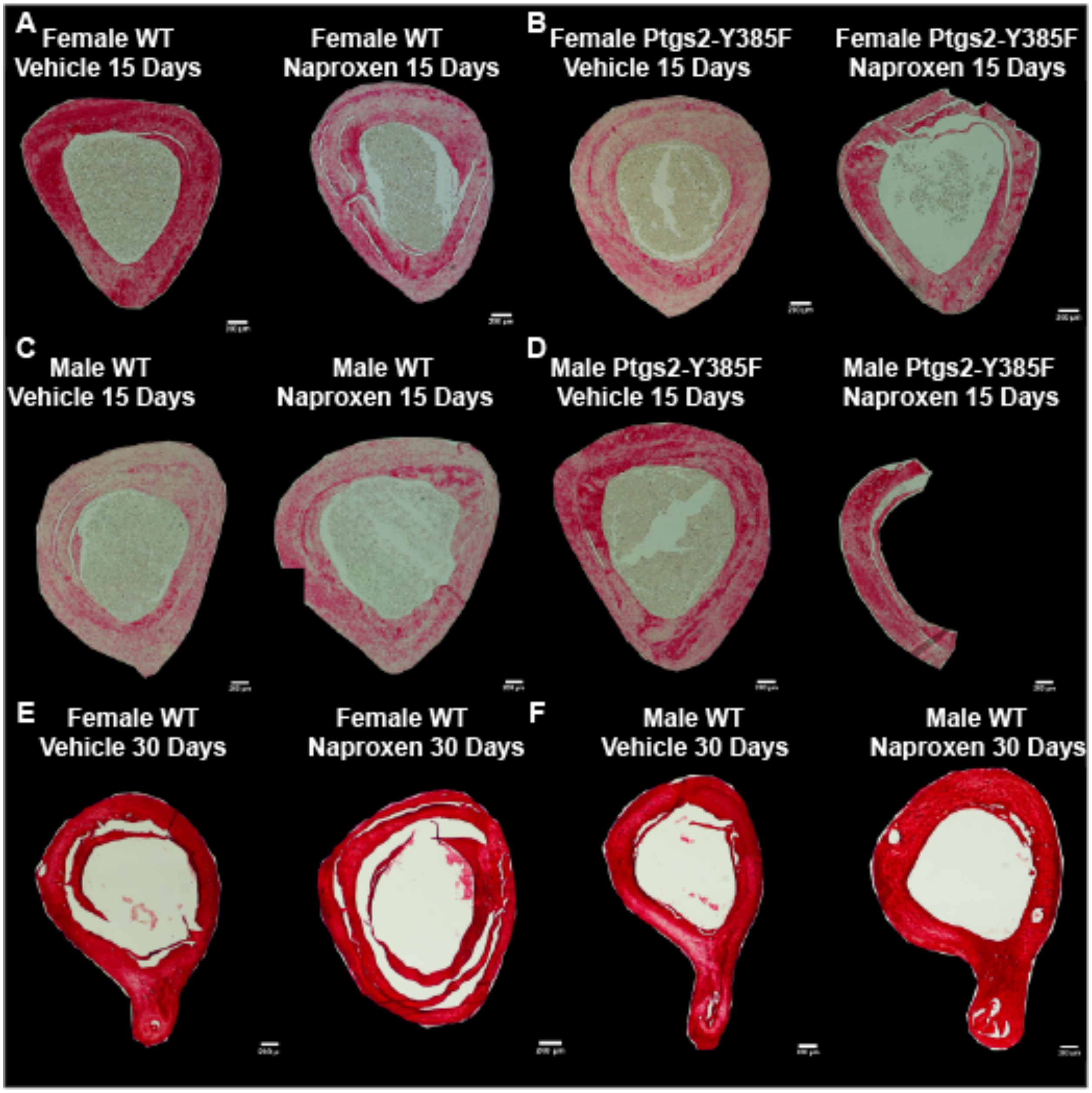


**Supplementary Figure 12: Representative bright field microscopy images of picrosirius red stained sections of tibias in WT and Ptgs2-Y385F mice who received naproxen or vehicle treatment for 15 or 30 days.** Quantifications of sections imaged under polarizing light are included in Figures 3 and 7 and Supplementary Figures 5 and 11. Brightfield images of picrosirius red staining were used to identify which technical replicate from each mouse was included in the analysis. Samples were chosen based on having the most evenly distributed red staining and minimal tissue folding.

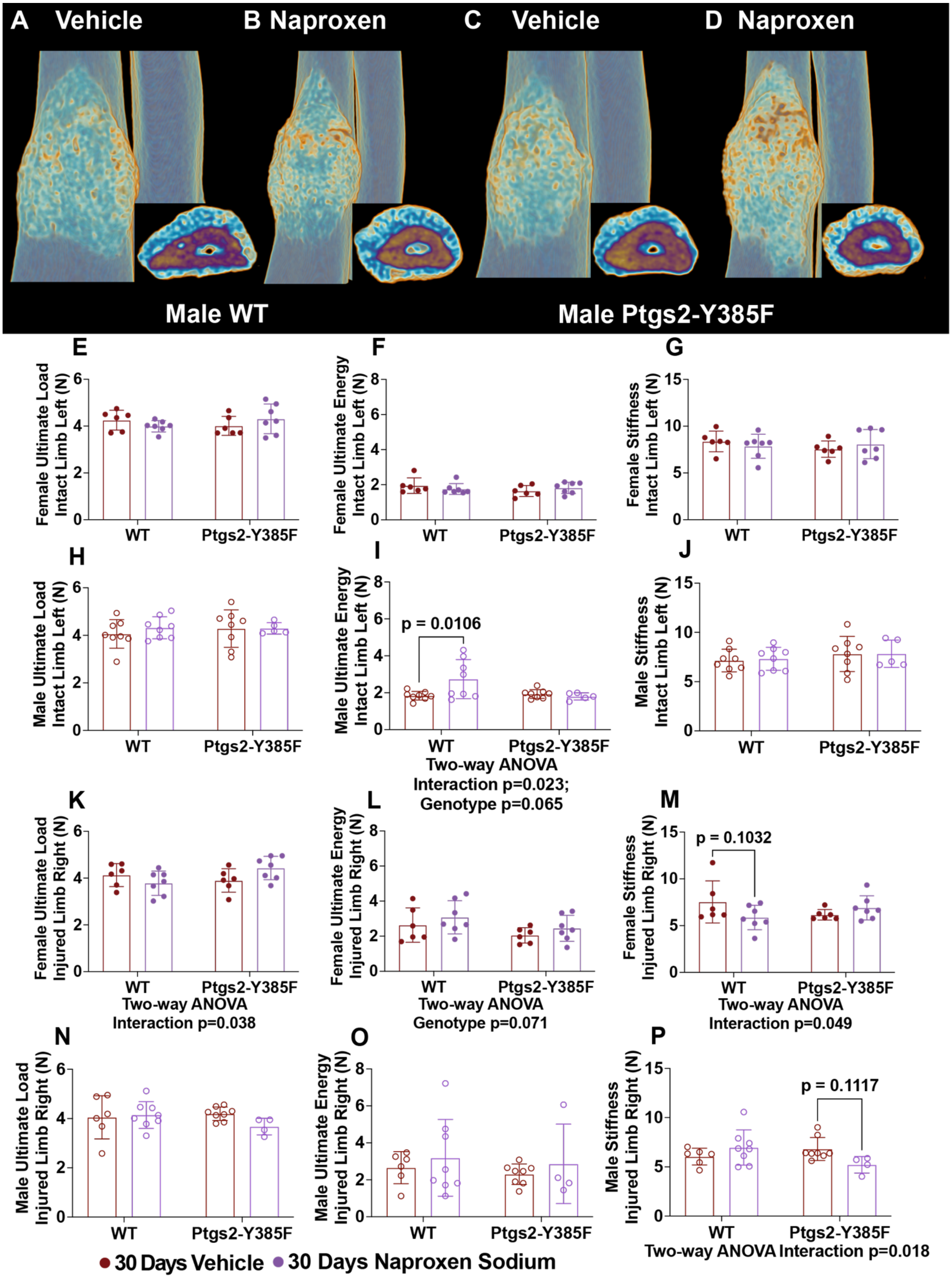

A Vehicle
B Naproxen
C Vehicle
D Naproxen
Male WT
Male Ptgs2-Y385F
E
Female Ultimate Load Intact Limb Left (N)
F
Female Ultimate Energy Intact Limb Left (N)
G
Female Stiffness Intact Limb Left (N)
H
Male Ultimate Load Intact Limb Left (N)
I
Male Ultimate Energy Intact Limb Left (N)
p = 0.0106
Two-way ANOVA Interaction p=0.023; Genotype p=0.065
J
Male Stiffness Intact Limb Left (N)
K
Female Ultimate Load Injured Limb Right (N)
Two-way ANOVA Interaction p=0.038
L
Female Ultimate Energy Injured Limb Right (N)
Two-way ANOVA Genotype p=0.071
M
Female Stiffness Injured Limb Right (N)
p = 0.1032
Two-way ANOVA Interaction p=0.049
N
Male Ultimate Load Injured Limb Right (N)
O
Male Ultimate Energy Injured Limb Right (N)
P
Male Stiffness Injured Limb Right (N)
p = 0.1117
Two-way ANOVA Interaction p=0.018
WT
Ptgs2-Y385F
30 Days Vehicle
30 Days Naproxen Sodium

**Supplementary Figure 13: Mice of both genotypes display sexually dimorphic effects of naproxen on fatigue fracture healing as measured by mechanical performance of the injured and intact limbs.** (A-D) Representative microCT reconstructions of male mouse forelimbs after 30 days of treatment showing fracture calluses and cross sections of the injuries approximately where the fracture crack begins. The mechanical performance of the bone callus was measured using monotonic compression to failure and the individual limbs were analyzed with sexes separated. (E) Calculated ultimate load, (F) ultimate energy, and stiffness of (G) female intact (left) limbs. (H-J) Corresponding measures for male intact limbs. (K) Calculated ultimate load, (L) ultimate energy, and stiffness of (M) female injured (right) limbs. (N-P) Corresponding measures for male intact limbs. n=6-7 females and n=5-9 males per group. A p value below 0.05 was considered significant and a p value below 0.1 was considered trending. Female replicates are displayed as filled circles and male replicates as empty circles.
